# Evidence for a selective link between cooperation and individual recognition

**DOI:** 10.1101/2021.09.07.459327

**Authors:** James P. Tumulty, Sara E. Miller, Steven M. Van Belleghem, Hannah I. Weller, Christopher M. Jernigan, Sierra Vincent, Regan J. Staudenraus, Andrew W. Legan, Timothy J. Polnaszek, Floria M. K. Uy, Alexander Walton, Michael J. Sheehan

## Abstract

The ability to recognize and discriminate among others is a frequent assumption of models of the evolution of cooperative behavior. At the same time, cooperative behavior has been proposed as a selective agent favoring the evolution of individual recognition abilities. While theory predicts that recognition and cooperation may co-evolve, data linking recognition abilities and cooperative behavior with fitness or evidence of selection are elusive. Here, we provide evidence of a fitness link between individual recognition and cooperation in the paper wasp *Polistes fuscatus*. Nest founding females in northern populations frequently form cooperative multiple foundress nests and possess highly variable facial patterns that mediate individual recognition. We describe a dearth of cooperative nesting, low phenotypic diversity, and a lack of individual recognition in southern populations. In a common garden experiment, northern co-foundress associations successfully reared offspring while all cooperative southern groups failed to rear any offspring, suggesting a fitness link between individual recognition and successful cooperation. Consistent with a selective link between individual recognition and cooperation, we find that rates of cooperative co-nesting correlate with identity-signaling color pattern diversity across the species’ range. Moreover, genomic evidence of recent positive selection on cognition loci likely to mediate individual recognition is substantially stronger in northern compared to southern *P. fuscatus* populations. Collectively, these data suggest that individual recognition and cooperative nesting behavior have co-evolved in *P. fuscatus* because recognition helps mediate conflict among co-nesting foundresses. This work provides evidence of a specific cognitive phenotype under selection because of social interactions, supporting the idea that social behavior can be a key driver of cognitive evolution.

## Introduction

The relationship between cognitive abilities and social structure is of long-standing interest to biologists. The social intelligence hypothesis (or social brain hypothesis) posits that selection pressures associated with social relationships in complex societies are an evolutionary driver of cognitive complexity^1–3^. For highly social animals, the abilities to track relationships within the group, cooperate with others, and predict how other individuals will behave in certain situations are considered to be cognitively challenging tasks that may impact individual fitness. Support for this hypothesis comes from comparative studies showing that cognitive performance^4–6^ and neuroanatomical proxies for cognition^1,7–9^ covary with proxies for social complexity, such as group size or mating system. Recently, general cognitive performance has been linked to group size and fitness in Australian magpies^10^. Additional indirect evidence of selection on cognition imposed by social systems comes from studies of brain gene expression showing, for example, shared transcriptomic signatures of monogamy across divergent vertebrate clades^11^. However, the evidence for the social intelligence hypothesis has come into question because predicted patterns do not hold for some clades and the use of different proxies for cognition and social complexity yields conflicting results^12–17^. More importantly, because of the reliance on such proxies, it has been difficult to identify specific cognitive traits that are under selection to facilitate social interactions.

Models for the evolution of cooperation frequently invoke animal recognition abilities as key mechanisms facilitating the evolution of cooperative behaviors^18–21^. Whereas kin recognition facilitates cooperation between relatives^22^, individual recognition has been identified as a building block of social cognition because it allows for cooperation between unrelated individuals^23^. Theory indicates that individual recognition enables cooperation because it allows for the identification of group members and reciprocity between individuals^18,19,24,25^. Indirect evidence of the fitness benefits of individual recognition for cooperative relationships comes from studies showing that territorial animals have higher reproductive success when they have familiar neighbors^26–28^. This result is presumably due to the decreased costs of conflict with territory neighbors when neighborhoods have stable compositions and established “dear enemy” relationships, in which they respect mutual territory boundaries and are less aggressive to each other than they are to strangers^29,30^. Whether individuals that do or do not recognize others vary in fitness outcomes in relation to cooperative behavior, however, has received less empirical attention. Overall, a major limitation to our understanding of the evolution of social cognition is direct evidence of a selective advantage of individual recognition in cooperative groups.

Here, we test the hypothesis that cooperative nesting selects for individual recognition in the northern paper wasp (*Polistes fuscatus*). This species provides an excellent study system for understanding the relationship between individual recognition and cooperation because both behaviors have been reported to vary across populations of this species^31,32^. Female *P. fuscatus* found nests in the spring, either as solitary foundresses or cooperatively with other foundresses. When females found nests cooperatively, they establish an aggression-based dominance hierarchy that determines the amounts of reproduction and work that each individual does^33–35^. Conflict among co-foundresses manifests in aggression between individuals and egg-eating^36^. Individual recognition has been hypothesized to function as a behavioral mechanism that maintains stable dominance hierarchies and minimizes conflict among co-foundresses^37^. The evolution of individual recognition in *P. fuscatus* is associated with increased phenotypic diversity due to the evolution of individually distinctive facial color patterns which function as identity signals and facilitate recognition^38,39^ as well as perceptual and cognitive mechanisms related to recognition^40,41^. However, a selective link between cooperation and individual recognition has yet to be demonstrated. Within-species variation in recognition and patterns of cooperation^31,32^ provides a powerful system to test for an evolutionary relationship between the two traits. In this paper, we test the hypothesis that cooperation selects for individual recognition using a combination of (1) common garden fitness assays of cooperative nesting behavior between populations with and without recognition, (2) an analysis of geographic clines in identity signaling and cooperation, and (3) population genomic analyses of the strength of selection on cognition loci. These three distinct lines of evidence are all consistent with an evolutionary scenario where cooperation among paper wasp co-foundresses has selected for individual recognition, an evolutionarily novel cognitive ability in northern *P. fuscatus* populations.

## Results

### Northern and southern populations of *Polistes fuscatus* differ in color pattern diversity and rates of cooperation

Northern populations of *Polistes fuscatus* in New York and Michigan have highly variable color patterns on their faces^38,39,42^; Fig. 1a), regularly form multi-foundress cooperative nests^43^, and have been shown experimentally to recognize individuals^42,44^. Here, we identified *P. fuscatus* populations at the southern end of the species range in Louisiana and coastal Georgia that have relatively invariant red faces and low levels of cooperative nesting (Fig. 1a, b). Using whole genome resequencing, we confirmed that these wasps are *P. fuscatus*. Wasps from these southern populations are closely related to those from northern populations, with low genetic differentiation between populations (F_ST_ = 0.07), matching previous findings of long-distance gene flow in *P. fuscatus*^45^. Further, wasps collected from across the range form a monophyletic clade, with wasps from southern populations interspersed with more northern populations (Fig. 1c and S1).

**Figure 1.**
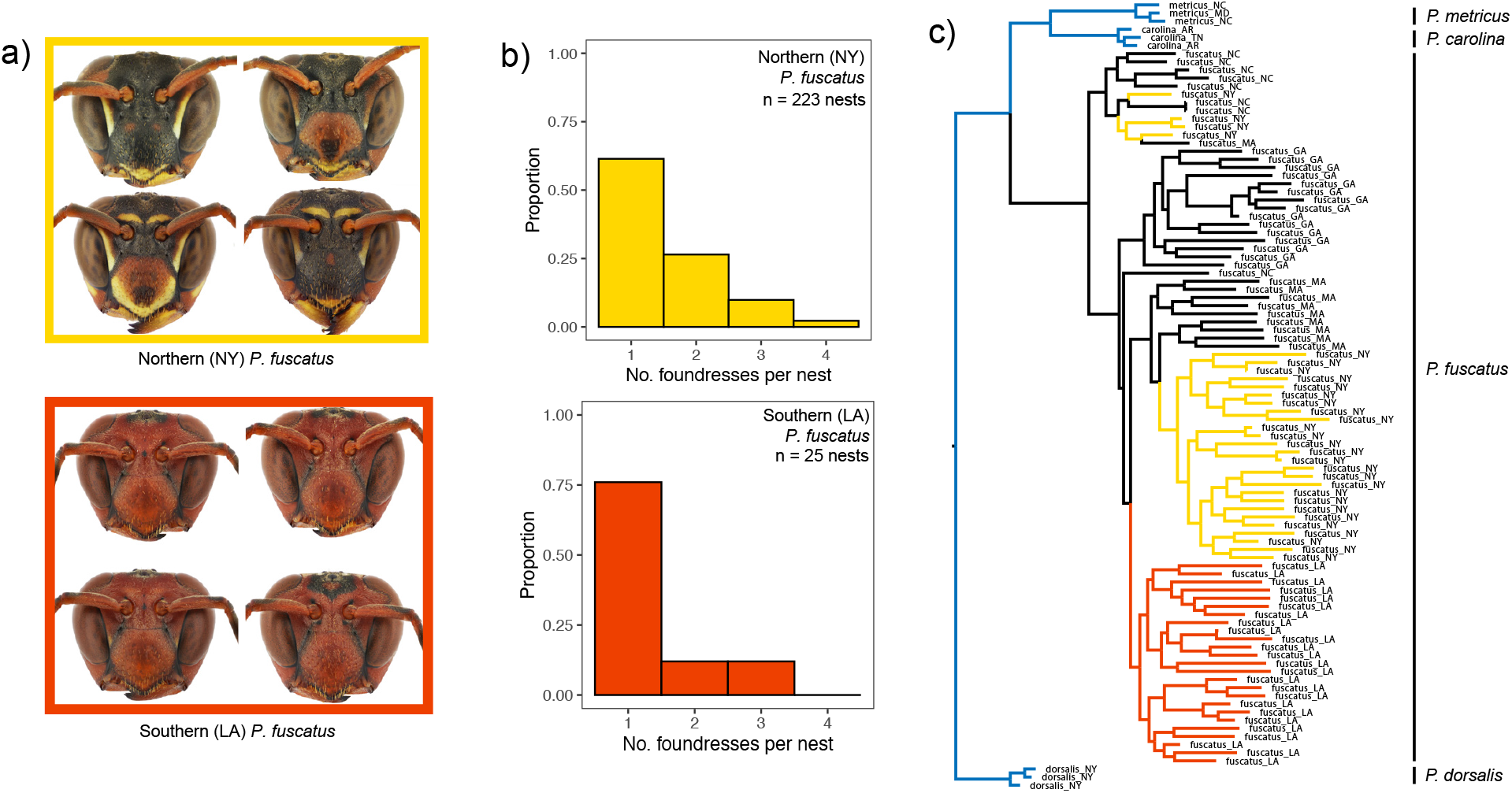
Northern and southern populations differ in face color pattern diversity and cooperation rates. a) Photographs of the faces of wasps from New York (northern; top) and Louisiana (southern; bottom), representing the diversity of face color patterns observed in these two populations. b) Histograms showing the distribution of the number of foundresses per nest in New York and Louisiana populations, demonstrating greater cooperation in New York. c) A phylogeny generated from SNP data from whole genome sequencing of *Polistes fuscatus* from across the geographic range, confirming that samples from northern and southern populations cluster together as a monophyletic clade, indicating they belong to the same species. Three closely related species (*P. metricus*, *P. carolina*, and *P. dorsalis*) are included as outgroups. Species name and US state of origin are given for each DNA sample. Branches are colored to highlight samples from New York (yellow) and Louisiana (red), samples from North Carolina, Massachusetts, and Georgia are black, and outgroups are colored in blue. See Fig. 5 for example faces of *P. fuscatus* from other populations.

### Populations differ in individual recognition behavior

Individual variation in color patterns of female *P. fuscatus* from New York and Michigan has previously been shown to mediate individual recognition^38,42^. Conversely, the lack of color pattern variation in other species of *Polistes* is associated with a lack of individual recognition^39^. Therefore, we reasoned that southern *P. fuscatus* populations lacking variable color patterning would also fail to show individual recognition. To test for individual recognition, we compared aggression between encounters of familiar and unfamiliar wasps from populations at the northern and southern portions of the range of *P. fuscatus*, following previous studies^32,39,44,46^. We compared the aggression between pairs of wasps that interacted for the first time (Day 0, ‘unfamiliar’) with their aggression when they met again two days later (Day 2, ‘familiar’). We controlled for the possibility of a general decrease in aggression over time that was not specific to a particular individual by also measuring the aggression between pairs of unfamiliar wasps on Day 1 and Day 3 of the experiment (*n* = 40 northern wasps, 42 southern wasps, 164 total trials). To compare the behavior of wasps from these two populations in the same experiment and at the same life stage, we collected foundress-destined female wasps (“gynes”) in the fall from northern and southern populations and overwintered them in the lab so they could emerge from overwintering at the same time in a controlled environment.

Wasps from both populations interacted most aggressively on the first day (Day 0) of the experiment (Fig. 2). There were significant differences in aggression between days for both northern (*χ*^2^ = 10.66, *p* = 0.014) and southern wasps (*χ*^2^ = 21.78, *p* < 0.001). However, only northern wasps showed evidence of individual recognition; northern wasps were significantly less aggressive to familiar individuals they encountered on Day 2 compared with the first time they encountered these individuals on Day 0 (*t* = 3.21, *p* = 0.010). However, they did not show significantly less aggression to other unfamiliar individuals on Day 1 (*t* = 1.79, *p* = 0.288) or Day 3 (*t* = 2.11, *p* = 0.159) relative to their initial aggression on Day 0. In contrast, southern wasps showed a general decrease in aggression after Day 0 regardless of whether the wasp they were interacting with was familiar or unfamiliar. Compared with Day 0, southern wasps were significantly less aggressive on Day 1 (*t* = 3.25, *p* = 0.009), Day 2 (*t* = 4.36, *p* < 0.001), and Day 3 (*t* = 3.45, *p* = 0.005). Overall, these data indicate that southern wasps use simpler decision rules and show a generalized decrease in aggression with repeated social interactions, while the decrease in aggression of northern wasps was specific to particular individuals they had encountered previously.

**Figure 2.**
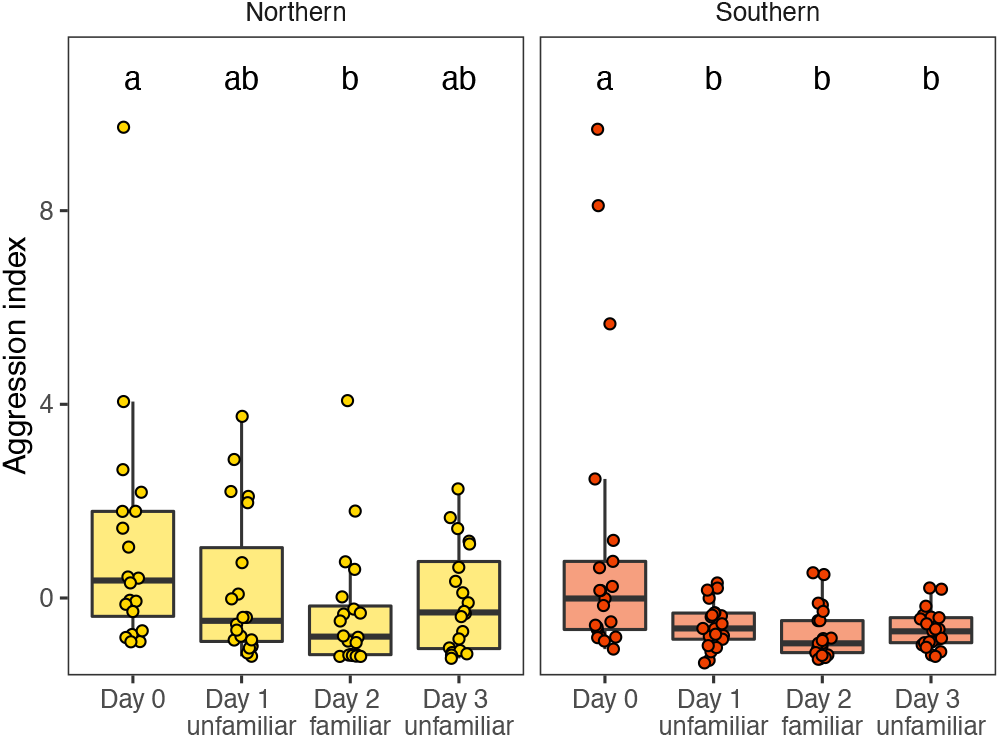
Results from experiments testing for individual recognition in northern and southern populations. On Day 2, wasps interacted with the same individual they interacted with on Day 0, while on Days 1 and 3, they interacted with individuals they had never encountered before. Different letters indicate days that are significantly different from each other from post-hoc comparisons within each population. Boxplots and individual data points show the aggression index computed from a PCA of the total numbers and durations of aggressive behaviors exhibited by both wasps during the trials.

### Recognition abilities are associated with differences in social organization between populations

The ability to recognize and discriminate among potential social partners is predicted to shape social networks and influence how animals interact with each other^47,48^. Therefore, we assessed whether recognition differences between northern and southern populations manifested in different social organizations and interactions. We established a lab common garden experiment in which the lab-overwintered foundresses were housed in groups of four individuals per enclosure. Each enclosure included four nesting huts, construction paper to provide nesting material, and *ad libitum* food and water. These groups were composed of three individuals from one nest of origin and another individual from a different nest. This design was chosen to reflect how cooperative nesting associations are often thought to occur, with co-foundresses often being former nestmates, but with unrelated foundresses sometimes joining nests^33,49^. We constructed social networks based on nocturnal associations of individuals before nests were established; paper wasps often “huddle” together in groups (or “cluster”, *sensu* West-Eberhard, 1969) when they are not on a nest. Therefore, we recorded which individuals were huddling together each night.

Social networks varied between enclosures of northern versus southern wasps. The mean huddle size per group was larger for southern wasps than northern wasps (*W* = 20, *p* = 0.006, Fig. 3a), and the within-group variance in huddle size was greater for southern wasps than northern wasps (*W* = 16, *p* = 0.003, Fig. 3b). These results suggest that southern wasps are more gregarious but form less stable associations than northern wasps. We tested this idea more directly by using these associations to construct social networks for each group. From these social networks, we computed what we define here as “edge evenness”. Analogous to species evenness in ecology^50^, edge evenness describes how evenly distributed relationships are across the network. Networks in which individuals interact at similar rates with all other individuals in the network have higher edge evenness than those in which some pairs or trios of individuals have stronger relationships than others. Social networks of southern wasps showed relatively even associations among individuals with little apparent sub-structure in the network (Fig. 3c, Fig. S3). In contrast, networks of northern wasps were characterized by stronger associations between pairs or trios of individuals to the exclusion of other individuals (Fig. 3c, Fig. S2). The edge evenness of southern wasps was greater than that of northern wasps (*W* = 19, *p* = 0.003, Fig. 3d). Overall, these data suggest that southern wasps are more gregarious but less discriminating about who they associate with.

**Figure 3.**
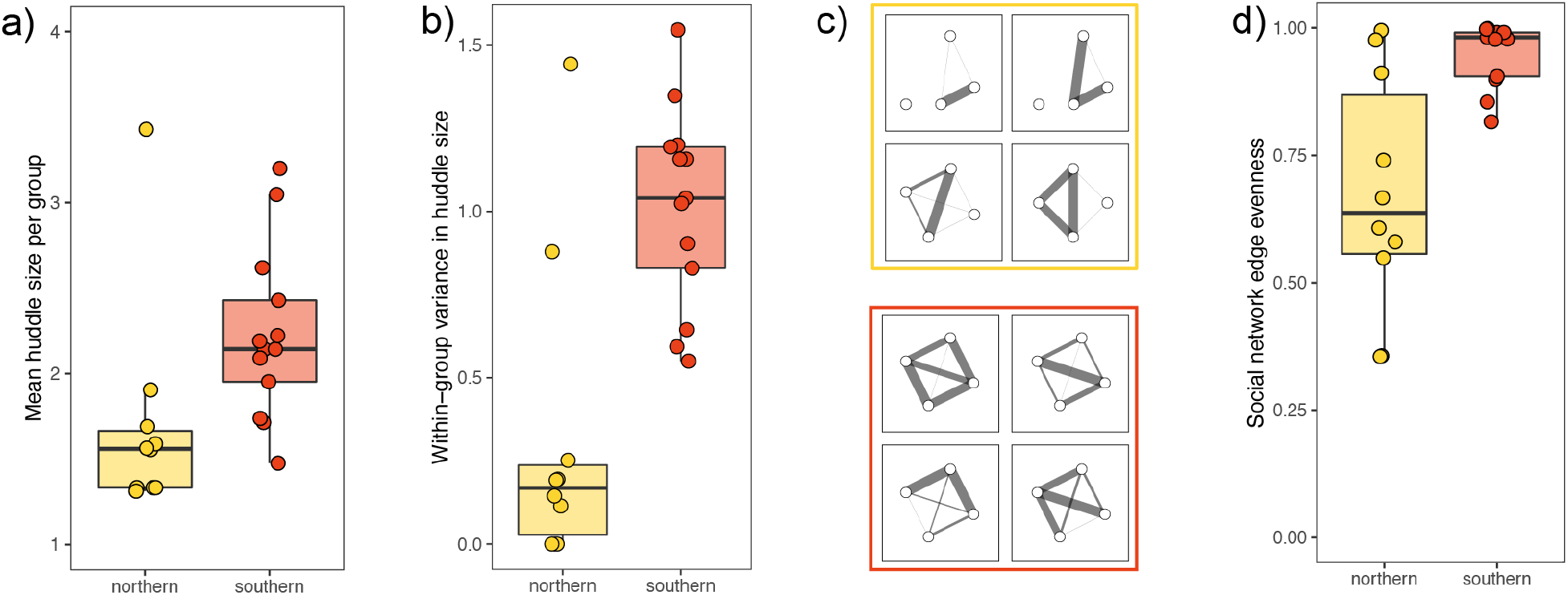
Pre-nesting associations of northern wasps are smaller, more stable, and less evenly distributed across the social network compared with those of southern wasps. a) Mean number of individuals per huddle (huddle size) per group prior to nest establishment. b) Within-group variance in huddle size across observation days. c) Representative social networks derived from pre-nesting association data for northern (top) and southern (bottom) wasps (networks for all groups can be found in supplemental figures S2 and S3). d) Edge evenness of social networks for northern and southern wasps. Higher values indicate connections are relatively evenly distributed among individuals in a network, while lower values indicate more skewed networks with stronger subgroups within the network.

### Cooperative southern nests are unsuccessful and fail to rear brood

Individual recognition in *P. fuscatus* is hypothesized to be an important behavioral mechanism stabilizing dominance hierarchies and reducing conflict among co-foundresses^37^. Therefore, we monitored the groups of wasps in the lab common garden experiment to compare the nesting success of multi-foundress wasps from both populations. Wasps from both populations started nests in the lab at similar rates. Among the groups that initiated nests in the lab, southern groups showed less stable nesting associations and less evidence of successful cooperation. Nests began with small pedicels attached to the ceilings of the cardboard huts and then were built out several cells at a time. In total, 4 northern groups and 3 southern groups tended nests. The nests were established between 4 and 12 days after housing. Interestingly, both northern and southern nests had multiple foundresses, and the mean number of foundresses per nest was similar between populations (Fig. 4a). However, the number of foundresses observed on a nest was not stable through time, and foundress number varied more for some nests than others (Fig. 4b). Overall, there was a trend of greater variance in the number of foundresses per nest for southern nests than northern nests (Fig. 4c), suggesting southern foundress associations were less stable.

**Figure 4.**
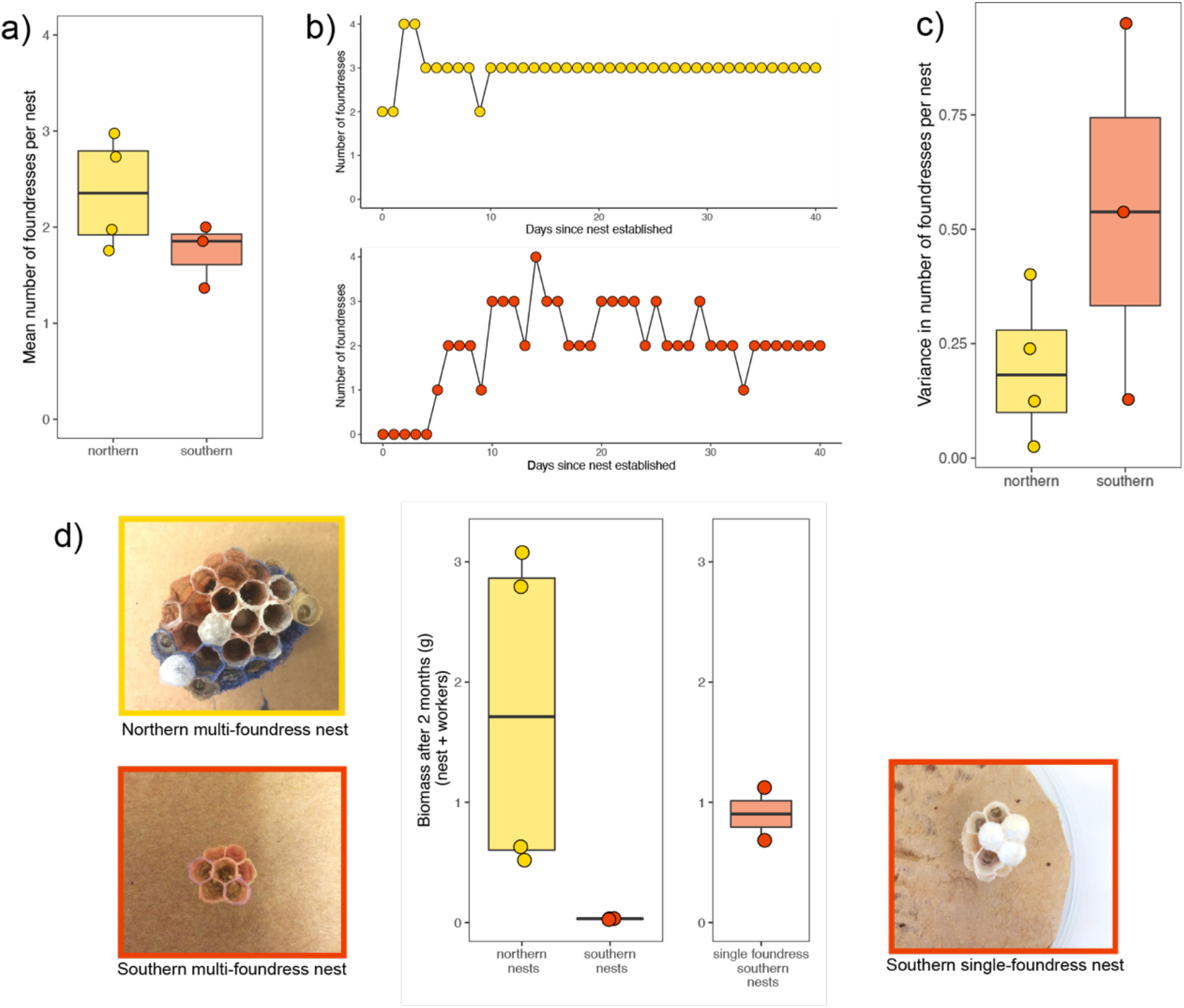
Southern multi-foundress nests failed to rear offspring. a) The mean number of foundresses per nest. b) The number of foundresses observed on a nest each day, for two example nests, displaying differences in the stability of foundress associations for different nests. c) Variance in the number of foundresses per nest. d) Nest biomass after two months of development, with photographs of example nests for each population. Also shown are nest biomass for southern single-foundress nests, which displayed normal development under the same conditions.

Strikingly, all the southern multi-foundress nests failed to develop; these nests were built to between 7 and 8 cells, and eggs were laid in these cells, but the eggs never developed into larvae and the nests were never expanded with additional cells. In fact, new eggs continued to be laid in cells throughout the two months of the experiment, clearly indicating previous eggs were eaten or removed by foundresses on the nest (Fig. 4d). In contrast, all the northern multi-foundress nests developed normally, with eggs developing into larvae and pupae (Fig. 4d). Cells continued to be added to these nests, and the number of cells after two months ranged from 17 to 31. Two northern nests had workers successfully emerge after two months. Overall, the biomass of northern nests was much greater than that of southern nests at the end of two months (Fig. 4d). Additionally, we know that the failure of southern multi-foundress nests to develop in our lab was not due to some problem with housing conditions that were specific to southern wasps, because two solitarily housed southern wasps built successful nests and reared offspring to pupation as single foundresses under the same lab conditions at the same time (Fig. 4d). This nesting experiment suggests that instability and conflict among southern co-foundresses prevented nesting success, providing evidence that individual recognition in northern populations is key to enabling stable dominance hierarchies and successful cooperation among co-foundresses.

### Identity signal diversity is associated with geographic variation in cooperation rates

Comparisons of recognition and nesting behavior between the northern and southern populations of *P. fuscatus* provide empirical support for the hypothesis that mediation of conflict and cooperation among co-nesting foundresses has been a selective agent favoring individual recognition in this species. If cooperative nesting has been a selective agent favoring the evolution of individual recognition in *P. fuscatus*, then identity signals that facilitate recognition should co-vary with rates of cooperative nesting across the species range, with regions with higher rates of cooperative nesting also showing greater color pattern diversity.

We collected female *P. fuscatus* across the East Coast of North America, spanning much of its latitudinal geographic range, and measured color pattern diversity (Fig. 5a). We chose to focus on the latitudinal gradient because of previous work suggesting cooperation rates in the north are higher for this species^31^. To quantify color pattern variation, we photographed faces and used a novel multi-step methodology for extracting homologous color patterns from images. We first normalized the lighting in the photos and aligned face images using homologous landmarks. We then segmented the images by color, by forcing each pixel to the nearest of three colors: yellow, red, or black (example images in Fig. S6). Finally, we subjected these color-segmented face images to a PCA and used a statistically significant set of 24 components to characterize variation among faces (Fig. S7, Table S2). We only sampled one wasp per nest to reduce the possibility that individuals in the sample were close relatives who might bias samples to being more homogeneous since color patterning is highly heritable in this species^51^. We computed the pairwise Euclidean distance between faces in PCA space for each site and took the mean of these distances as our measure of face diversity for a site. There was a strong positive relationship between latitude and face diversity in a site (*R*^2^ = 0.74, *F*_1,16_ = 46.1, p < 0.001; Fig. 5b). The relative lack of facial diversity was especially pronounced in the southernmost populations from Louisiana and coastal Georgia, which occur below 32° latitude (Fig. 5b). Compared with these southernmost populations, face diversity was about 1.6 times higher at around 35° latitude in South and North Carolina, with diversity increasing further in more northern populations (Fig. 5b).

**Figure 5.**
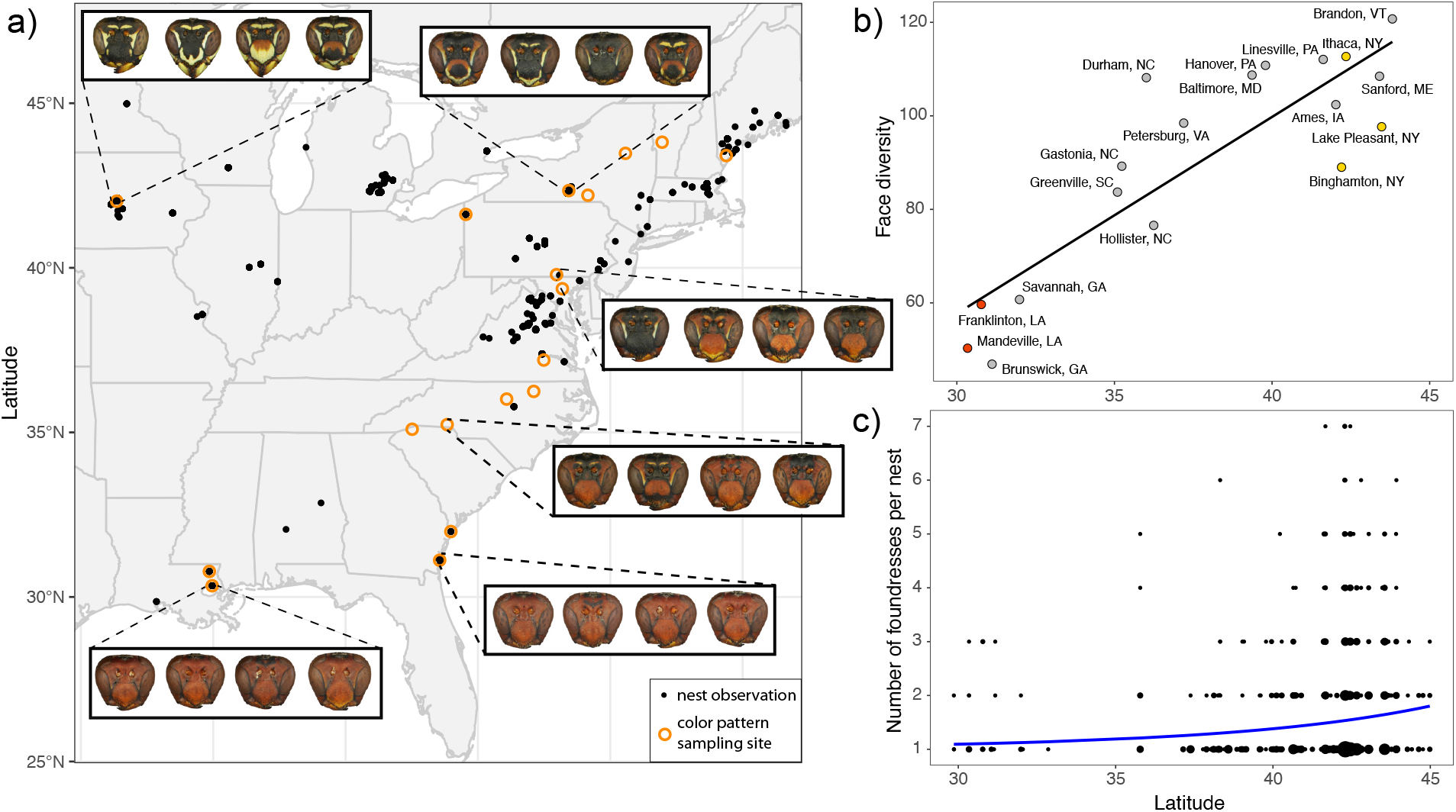
Color pattern diversity and cooperative nesting rates both increase with latitude. a) Map of sampling locations for color pattern diversity and cooperation rates of *P. fuscatus* wasps. Black points represent nest observations taken from the WASPnest dataset^31,43^ as well as new observations reported in this paper. Orange open circles mark sites where we collected and photographed wasps to measure color pattern variation. Also shown are photographs of representative individuals from several sites to demonstrate the color pattern variation across the range. b) The relationship between face diversity and latitude across the range of sampling sites fit with a linear regression line. Face diversity was measured as the mean distance between faces within a population in PCA scores computed from color segmented images. Points representing sites in Louisiana are colored red, and those representing sites in New York are colored yellow. c) The relationship between the number of foundresses per nest and latitude fit with a zero-truncated Poisson regression line. The sizes of points are scaled according to the number of observations

We observed nesting behavior in southern wasp populations and added these data to previously published datasets of nesting behavior in *P. fuscatus*^31,43^. There is a positive relationship between the number of foundresses per nest and latitude (z = 6.81, p < 0.001, *n* = 2,021 nests, Fig. 5c), consistent with the findings of earlier studies^31^. At the southern end of the range, the majority of foundresses nest solitarily, whereas at the northern end of the range, most foundresses are part of cooperative groups (e.g. 56% solitary in Louisiana, 60% cooperative in New York). Additionally, the occasional cooperative nests that were observed in the southern portion of the range never had more than 3 foundresses. At northern latitudes, large nesting associations of 4 or 5 foundresses occur with some regularity, and groups of 6 or 7 foundresses are observed as well (Fig. 5c). Results from these two clinal datasets are consistent with the hypothesis that cooperation selects for individual recognition by favoring individuals who signal their identity.

### Genomic evidence of selection on cognition associated with individual recognition

Previous population genomics studies of northern *P. fuscatus* populations identified multiple strong recent selective sweeps in genomic regions related to learning, memory, and visual processing^52^. Evidence of selection on learning, memory, and visual processing was substantially weaker in two closely related species, *P. metricus* and *P. dorsalis*, which lack individual recognition, suggesting that the patterns of selection in northern *P. fuscatus* populations may be associated with individual recognition. If the signatures of selection on these loci are the result of an evolutionary advantage of individual recognition in the more cooperative northern populations, these same loci are predicted to show weaker or no evidence of selection in southern populations due to the absence of individual recognition. Therefore, we repeated this analysis using southern populations to directly compare evidence of recent selective sweeps between northern and southern populations. Selective sweeps were identified using SweepFinder2^53^, which uses deviations in the local site frequency spectrum to generate a composite likelihood ratio (CLR) of a selective sweep in that genomic region. Larger CLR values provide evidence of stronger selection, more recent selection, selection on newer mutations, or some combination of these phenomena^52^.

Both northern and southern populations show evidence of recent strong positive selection, with some selective sweeps shared across populations and other selective sweeps that are unique to only one population (Fig. 6a, S8). We assessed patterns of selection on loci that likely contribute to cognitive abilities underlying individual recognition using two approaches. First, we compared scaled CLR values between northern and southern populations for loci annotated with gene ontology (GO) terms related to learning, memory, and visual processing, directly replicating the previously published analysis of northern *P. fuscatus* populations^52^. Scaled CLR values for these annotated “visual cognition genes” were elevated in both populations, but there was a significant interaction between population and gene type (gene type: χ^2^ = 82.43, *p* < 0.001; population: χ^2^ = 268.73, *p* < 0.001; gene type × population: χ^2^ = 28.50, *p* < 0.001), indicating that selection on visual cognition genes is stronger in northern populations than southern populations (Fig. 6b). Second, we compared scaled CLR values between northern and southern populations for genes that are differentially expressed during social interactions in northern *P. fuscatus*^54^. Experimental evidence for differential regulation in response to social interactions suggests these genes could play a role in recognition behavior in this species. Again, we find evidence of stronger selection on socially regulated genes in northern compared to southern populations (gene type: χ^2^ = 206.56, *p* < 0.001; population: χ^2^ = 78.85, *p* < 0.001; gene type × population: χ^2^ = 78.69, *p* < 0.001; Fig. 6c). Rather than comparing the relative evidence of selection across all genes, we can also ask whether genes in these two datasets are overrepresented among the most strongly selected genes. We find greater enrichment of strongly selected genes in northern compared to southern populations for both GO term and socially regulated gene sets (Table S3). Together, these data show that, compared with southern populations, selection in the north has been stronger on genes that are likely involved in the perceptual and cognitive abilities of wasps to recognize individuals and mediate social interactions.

**Figure 6:**
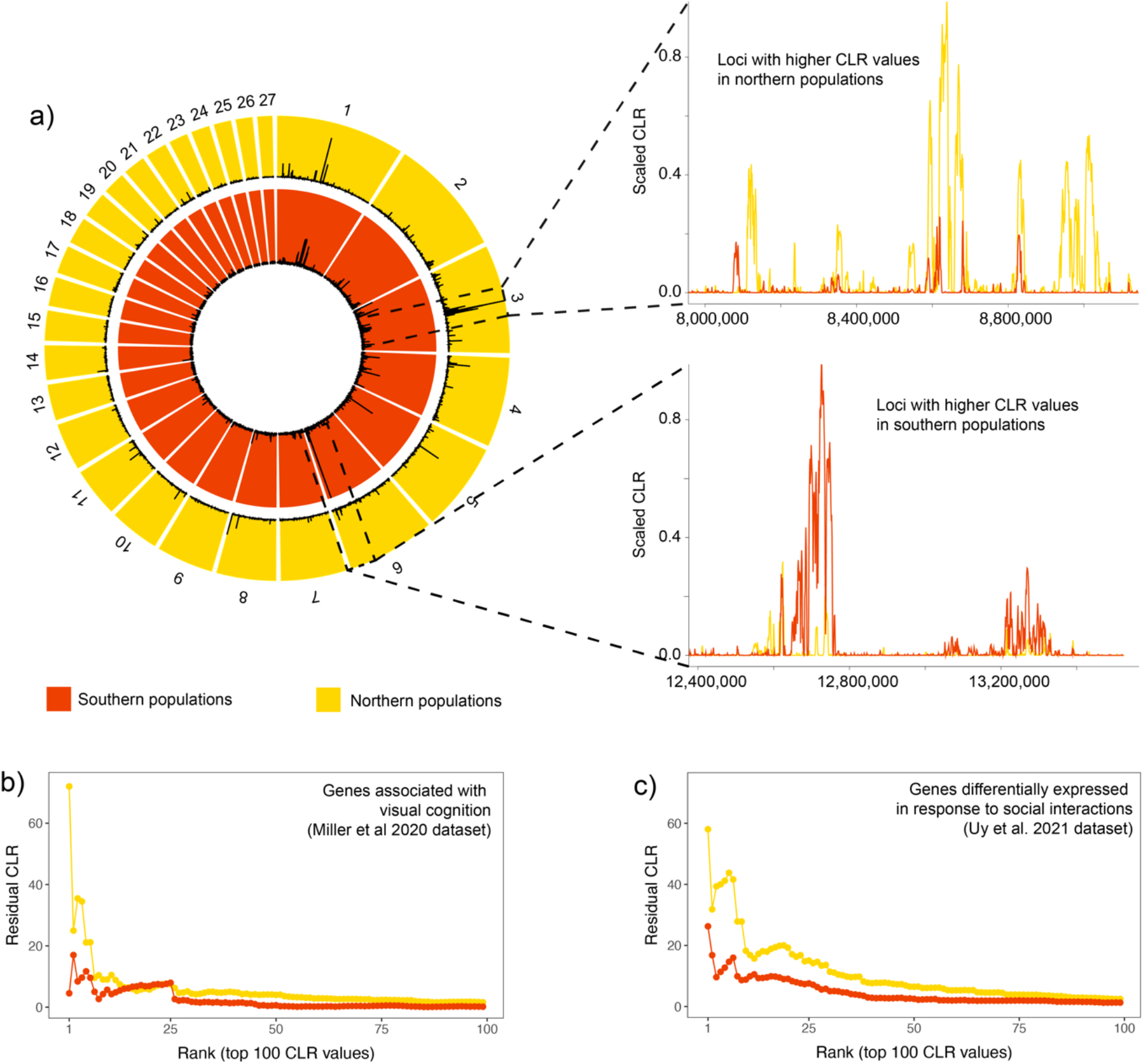
Stronger and more recent selection on candidate cognition loci in northern populations. a) Comparison of scaled composite likelihood ratio (CLR) values between northern (outer) and southern (inner) populations for the largest 27 scaffolds in the *P. fuscatus* genome. CLR values have been smoothed over 10,000 bp windows. Examples of regions where CLR values are greater in the north (top) and south (bottom) are shown. b & c) Residual CLR values for the top 100 CLR values of putative social cognition genes from datasets of a) genes with gene ontology (GO) terms related to learning, memory, and visual processing and b) genes that are differentially expressed in response to social interactions in northern populations. Both data sets show that northern populations have elevated signatures of selection on putative social cognition genes.

## Discussion

Despite longstanding interest in the evolutionary relationship between social organization and recognition, direct evidence that recognition abilities are under selection for social interactions has been missing. This limitation is due, in part, to the fact that within-species variation in recognition abilities has rarely been documented. Using within-species variation in cooperation and individual recognition in the northern paper wasp (*Polistes fuscatus*), we provide three distinct lines of evidence that together provide strong support for the hypothesis that cooperation selects for individual recognition. First, in a common garden lab nesting experiment, southern wasps lacking individual recognition (Figs 1–2) were more gregarious but had less stable social networks. Southern multi-foundress nests failed to successfully rear offspring, providing evidence of a direct fitness consequence of the absence of individual recognition in cooperative groups (Figs 3–4). Second, across the range of this species, color pattern diversity co-varies with rates of cooperation such that identity signals used for individual recognition are most apparent in the more cooperative northern populations (Fig 5). This pattern is consistent with expectations of selection favoring signalers who signal their identity to facilitate recognition in cooperative populations^38,55–57^. Third, genomic analyses reveal stronger signatures of selection in northern populations, compared with southern populations, on genes related to visual cognition and social experience (Fig 6). This pattern is consistent with expectations of selection favoring improved perceptual and cognitive abilities to recognize individuals in more cooperative populations^23,58,59^.

### Evolution of recognition abilities

The adaptive value of recognition and discrimination is often conceptualized as the shifting of optimal thresholds for “accepting” or “rejecting” individuals^60–62^. There are many examples of plastic shifts in acceptance thresholds showing that animals often adjust decision rules in an adaptive way^e.g.,63–66^, but less work has documented evolutionary changes in the decision rules that determine acceptance thresholds^67^. The results of our behavioral experiments suggest that southern *P. fuscatus* foundresses universally reject or accept conspecific foundresses depending on the context. Wasps from these populations are characterized by a lack of individual recognition, a generalized decrease in aggression to conspecifics through time, and gregarious, indiscriminate huddling behavior in pre-nesting associations. Similar patterns of aggression to southern *P. fuscatus* were seen in the closely related (and also rarely cooperatively nesting) *P. metricus* using the same trial design^39^. Thus, decision rules of southern *P. fuscatus* likely represent the ancestral state for this species. Therefore, the evolution of individual recognition in *P. fuscatus* is associated with increasing the specificity of acceptance and rejection decision rules depending on specific individuals rather than behavioral context, as evidence of discriminating behavior was observed in all behavioral experiments. The result that individual recognition in wasps evolved from an ancestral state of context-dependent universal acceptance provides an important example of how the decision rules guiding acceptance thresholds may be targets of selection during the evolution of recognition systems.

Our results have multiple implications for how social complexity relates to the cognitive demands of social life^68–70^. First, our results add to a growing literature demonstrating that group size and social complexity are not the same^9,70–72^. Initial expectations might be that individual recognition should be associated with larger social groups in general. This pattern is observed in the finding that the number of foundresses per nest and signal identity information covary latitudinally in *P. fuscatus* (Fig. 1b; Fig. 5c). However, southern wasps actually formed larger huddles, on average, but these huddles were less stable. Social network analysis revealed that northern wasps had stronger relationships among sub-sets of individuals to the exclusion of others, while southern wasps had relatively evenly distributed relationships across the network. This result is consistent with the idea that individual recognition allows for relational social complexity within groups^48,71^ and highlights that group size alone may be a poor proxy for social complexity in many contexts. Second, our work provides insights into which features of social relationships may drive increased cognitive complexity. Social interactions can involve cooperation and/or conflict, and both have been hypothesized to be cognitively demanding^17,69,73–76^. The common garden nesting experiments suggest that conflict among co-nesting foundresses is a main driver of selection for individual recognition. The southern wasps failed to rear offspring because of oophagy, a sign of conflict among the foundresses. These data argue for a role of recognition in facilitating cooperation by managing conflict.

Results from the common garden experiments shed light on the behavioral mechanisms favoring recognition but are only one means to test for evidence of selection linking cooperation and individual recognition. The results of our geographic sampling of color pattern and cooperation are consistent with expectations of selection favoring individuals who signal their identity to facilitate recognition in cooperative populations^38,55–57^. The extensive variation in color patterns within and between populations of *P. fuscatus* has long been a source of consternation and puzzlement for students of paper wasps^77,78^. Geographic variation in color patterning is commonly reported in insects and other animals and is frequently linked to selection imposed by the abiotic environment, predation, or sexual selection^e.g.,79–84^. Our data suggest social selection among female foundresses is the driver of color pattern variation in *P. fuscatus*. These data add to a growing body of research showing that identity information in signals often correlates with measures of social complexity, suggesting social environments can impose selection on signals to make individuals more recognizable^85–90^.

### Evidence for recent evolution of individual recognition in *P. fuscatus*

Individual recognition appears to be evolutionarily derived and unique to *P. fuscatus* among closely related species^39,91^. Further, population genomic analyses have revealed multiple selective sweeps within the last few thousand years that are enriched for genes likely involved in individual recognition, such as genes related to visual processing, cognition, learning, and memory^52^. Many of these selective sweeps occurred since the last glacial maximum when the Laurentide Ice Sheet covered much of the current northern range of *P. fuscatus*^92^. Together with our results demonstrating individual recognition and identity signals are absent in southern populations (Figs 2 and 5), these studies suggest a hypothesis in which ancestral populations lacking identity signals with low rates of cooperation recently evolved individual recognition as an adaptation to enable successful cooperation as the species expanded northward following the last glacial retreat. The ecological factors that favor cooperation at northern latitudes are currently unknown, but cooperative nesting decreases the probability of nest failure before workers emerge^34^, and shorter summers in northern climates might reduce the probability that solitary foundresses can make multiple nesting attempts and still succeed. It will be important to test this hypothesis in the future.

### Why do southern populations lack individual recognition?

Given the low population genetic structure at the continental scale of *P. fuscatus*^45^, population differences in color patterning and selection on social cognition suggest multiple possibilities for why we do not observe individual recognition or color pattern diversity in southern populations. First, it may be the case that alleles related to individual recognition arose recently in northern populations and have yet to reach southern populations. Evidence for this scenario comes from a previous analysis of selection in this species that demonstrated that many selective sweeps involved recent *de novo* mutations^52^. However, the lack of population structure suggests that the recent evolution of individual recognition is unlikely to fully explain the geographic pattern of coloration and recognition abilities, as we would expect recognition-associated alleles to quickly spread if they were beneficial in all populations. Indeed, migration of alleles under strong selection in northern populations into southern populations may explain some, though not all, of the shared signatures of selection found here. Another possibility is that individual recognition is costly in *P. fuscatus*, meaning it is only favorable when rates of cooperation are sufficiently high to make the benefits of recognition outweigh these costs. In particular, the cognitive abilities related to recognition are assumed to be costly in terms of growth and maintenance of the requisite neural tissues^93–95^. Low rates of cooperation in southern populations may then remove the potential benefits of the cognitive mechanisms related to individual recognition, so the alleles for these traits are selected against. Lack of recognition behavior would then also remove benefits of signaling identity via distinctive color patterns. A similar model may explain the lack of individual recognition described in a *P. fuscatus* population in mountainous regions of central Pennsylvania with relatively low rates of cooperation and relatively low color pattern diversity^32^. However, models of identity signal evolution suggest that increased signal diversity may be favored even under very small fitness benefits provided the costs of distinctiveness are very small or non-existent^56^. Thus, the absence of color pattern diversity in the southern populations suggests that there may be selection either against particular color pattern variants involved in identity signaling or selection favoring the red facial color pattern that is common throughout the Gulf coast region. Future comparative analyses of clinal variation in alleles associated with cognition and color patterning will be useful to help discriminating among the hypotheses raised by the present dataset.

### Conclusions

Social structure and cognitive abilities vary widely among animals. The extent to which they are linked has been an ongoing subject of debate, often involving proxies of both social behavior and cognition. Using three distinct types of studies examining common garden fitness assays, geographic patterns of behavior and signal diversity, and population genomic analyses of selection on cognition loci, we provide cohesive evidence that cooperation favors the evolution of individual recognition. Individual recognition is a bedrock of many complex social behaviors. Our study demonstrates that understanding the factors that shape the evolution of specific cognitive abilities rather than just brain size or other proxies of general cognition can provide clear evidence for a link between social behavior and cognitive evolution.

## Methods

### Individual recognition experiment

Experiments were performed on lab overwintered *P. fuscatus* gynes that were collected in the fall of 2019, from Northern (NY and ME) and southern (LA) populations. Individuals were overwintered in plastic deli cups along with their nestmates, and provided water and sugar, as well as crumpled construction paper in which to hide. They were overwintered for approximately three months at 4°C for northern wasps and 10°C for southern wasps, to account for natural differences in winter temperatures between these populations. Following overwintering, wasps were weighed, paint-marked on their thorax (Testors enamel paint), and housed individually in deli cups for 5-6 days before the start of the experiment at a temperature of approximately 23°C with 12:12 light–dark cycle.

Separately for each population, we ranked individuals by weight to create three weight classes of similarly sized individuals. We then paired individuals together such that they always encountered other individuals from different nests but from the same weight class. These criteria resulted in 40 northern and 42 southern wasps for the experiment. On Day 0 of the experiment, pairs of wasps were placed together in plastic petri dishes and their interactions were filmed for 45 mins. Immediately following this trial, the pair was housed together a new deli cup overnight to give the individuals additional time to become familiar with each other. Between 9 and 10 AM the next morning (Day 1) these paired wasps were then put into solitary housing where they remained for the rest of the experiment. On Day 1 and 3 of the experiment, wasps were paired and filmed interacting as described above but with new individuals they had never encountered before. On Day 2 of the experiment, they were paired again with the same individual they interacted with on Day 0. We additionally controlled for potential day effects by starting the experiment for half of the wasps on one day and the other half on the subsequent day. All interaction trials occurred during the afternoon (13:00-18:00) at temperatures ranging from 25 to 26°C.

We scored aggressive behaviors for the first 15 minutes of each trial using BORIS^96^. Our ethogram was developed based on a combination of established ethograms for *Polistes*^35^, and our own preliminary observations of the aggressive behaviors that are common in this type of experiment. We scored the following as point behaviors (instantaneous behaviors that are counted for each occurrence): dart, a rapid forward movement towards another individual; snap, open mandibles towards another individual; bite, mandibles closing on another individual; kick, rapid leg extension that appeared to push off or push away another individual. We scored the following as state behaviors (behaviors that have durations) and denoted the start and stop times: approach, orienting and moving towards another wasp to engage in an interaction; chase, one wasp pursuing another wasp who appears to be avoiding the interaction; antennation, probing another individual with antennae; grapple, wrestling-type behaviors with both individuals engaged with biting and kicking. Each behavior was coded to one of the two subjects. Observers were blind to treatments and experiment day when scoring behaviors.

For each trial (*n* = 164), we summed the total numbers of point behaviors, and summed the durations of all state behaviors. We included all behaviors associated with aggressive physical interactions (approach, bite, dart, dodge, kick, snap, antennation, chase, and grapple) in a principal components analysis (centered and scaled) using the ‘prcomp’ function in R. We took the first principal component, which explained 33% of the variation, as an aggression index (see Table S1 for factor loadings). For statistical models, the aggression index was log-transformed to better meet assumptions of parametric tests. Separately for each population, we fit linear mixed effects models of the aggression index using the *lme4* package^97^, with experiment day as a fixed effect and cohort as a random effect. Tukey adjusted post-hoc comparisons among experiment days were performed using the *emmeans* package^98^.

### Common garden lab nesting experiment

Lab overwintered wasps from the recognition trials were individually marked and housed in groups of four individuals: three individuals from one nest of origin and another individual from a different nest. Groups of wasps were housed in enclosures consisting of two 36.8 cm × 22.2 cm × 24.8 plastic Kritter Keepers (Lee’s Aquarium & Pet Products) stacked on top of each other, with ventilation holes drilled into the sides and top. Four 10 cm x 10 cm cardboard nesting “huts” were attached to the top of the enclosure to provide each wasp the option to either nest alone or co-found a nest with other individuals. Each enclosure was provided with ample crumpled cardboard paper to provide nesting material, as well as a sugar cube, honey, water, and, once nests were established, an *ad libitum* variety of larval insects (waxworms (*Galleria mellonella*), hornworms (*Manduca sexta*), and mealworms (*Tenebrio molitor*); Rainbow Mealworms). Wasps were kept in a temperature-controlled room under conditions meant to mimic warm summertime environments to stimulate nesting (14:10 light-dark cycle, 25-28°C daytime temperature, 21-25°C nighttime temperature, 20-40% humidity).

Before the lights came on each morning, we recorded the location of each individual relative to other individuals in the group as either: alone – greater than one body length from any other individual; in proximity – within one body length of another individual; or huddled – touching or close enough to be capable of touching another individual. Once a nest was established in an enclosure, we also recorded which individuals were on or next to the nest overnight for the duration of the experiment. Individuals often leave the nest to forage or acquire nesting materials during the day but return and remain on the nest at night^31,35^. Therefore, nighttime surveys provide a reliable measure of which individuals are associated with the nest. During these surveys we also visually inspected nests, counted cells, and recorded the most advanced larval stage observed in a nest. In total, 8 groups started nests, but one nest was quickly abandoned after only one day and is not included in nest descriptive statistics. We measured nest development by weighing all nests two months after housing.

We analyzed pre-nesting associations for the first two weeks of the experiment because all nests were established by two weeks into the experiment. For groups that did not build a nest, we used the full two weeks of data. For groups that built a nest, we only used data from before the nest was established. Similarly, 4 individuals from 4 different groups died during the experiment, so for these groups we also only used data from before one individual in the group died. Note, conclusions did not change when restricting the data of all groups to observations that occurred before any nests or deaths (first 4 days of the experiment). To compute descriptive statistics of the number of individuals per huddle (huddle size), we first computed the mean huddle size per group-per day, and then used these numbers to compute grand mean and variance for each group. We statistically compared the mean and variance in huddle size using Wilcoxon rank sum tests. Because of the small sample size of numbers of nests, we only report descriptive statistics of foundress associations and nest development.

### Social network analysis

We used the pre-nesting huddle data (above) to construct social networks for each group. Connections between individuals (“edges”) were weighted depending on whether individuals were huddled together (weight = 2) or simply in proximity (weight = 1). From these social networks, we computed what we define here as “edge evenness”, which is analogous to the species evenness metric in ecology, derived from the Shannon diversity index^50^. Edge evenness (*J’*) was computed as

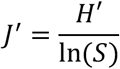

where *S* is the number of possible edges in the network, in our case 6 for a 4-individual network, and *H′* is the Shannon diversity index

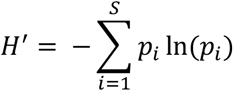

where *p_i_* is the proportion of weight of the *i*th edge in the network relative to the sum of all weights in the network. Edge evenness describes how evenly distributed edge weights are across the network. Networks in which individuals interact at the same rates with all other individuals in the network have an edge evenness of 1, while lower values indicate skewed networks in which some pairs or trios of individuals have stronger relationships than others. We statistically compared edge evenness between populations using Wilcoxon rank sum tests.

### Photography and color pattern measurement

Individuals were captured using nets, freeze-killed, and stored in a −20°C freezer for preservation. To photograph faces, we first removed the head and the antennae to allow full view of the color pattern. We photographed faces under standardized lighting conditions in the lab in a photographic tent using a Canon 6D camera and Canon 100mm macro lens. We confirmed that *P. fuscatus* faces do not reflect light in the ultraviolet range (Fig. S4), therefore standard camera equipment captures the full range of color variation in this species. Specimens were illuminated with bright, diffuse light to minimize shadows and glare by positioning three lights (compact fluorescent) facing away from the specimen to reflect off the walls of the photographic tent and surrounding the specimen with a cylinder of translucent plastic. To control for potential slight differences in lighting across days, we also photographed three spectrally flat gray standards (90%, 27%, and 3% reflectance: Color-aid gray set) under identical conditions during each photography session^99^.

Although there is some minor variation in brightness and hue within colors, it is clear that the meaningful variation among individuals occurs in patterns of black, red/brown, and yellow (Fig. 1, 5, S6). These three colors are present in most populations of this species and are also the primary colors observed across species of *Polistes*. Therefore, our goal in this analysis was not to measure color *per se*, but to objectively quantify color pattern and compare patterns in homologous regions across individuals. To do so, we first used the MICA toolbox^99^ to normalize the light levels across photographs using the gray standards photographed during each session. We then converted these normalized and linearized images using a CIE XYZ cone catch model that was specific to our camera and photography illuminant (Fig. S5) using the chart-based cone-catch model procedure in the MICA toolbox. We exported these images as .jpg files and adjusted the maximum pixel value to 0.4 out of 1 to make the image appear bright on the screen but without any pixel values being oversaturated.

We then used the R packages *patternize*^100^ and *recolorize*^101^ to align images, map color patterns, and analyze variation. First, we added 8 landmarks to each face image and then used the ‘alignLan’ function in *patternize* to align all of the images by these landmarks and mask areas of the image that fell outside of the main regions of interest, encompassing the clypeus, inner eye region, and frons (see Fig. S6). Then, we used *recolorize* to classify pixels in these masked images to three color clusters: black, red, and yellow. To do so, we first obtained a color palette by running an initial color segmentation step on a subset of 30 images that appeared representative of these three colors using the ‘histogram’ method with 6 bins per color channel using the ‘recolorize’ function and then implementing the ‘recluster’ function using a similarity cutoff of 15%. These parameters were chosen based on trial and error to create color segmented images that appeared similar to the color patterns in the original images. We clustered the colors by similarity to three color clusters and took the weighted average of these three clusters which resulted in a color palette corresponding to the black, red/brown, and yellow present in the images. We created a separate color palette for the southernmost populations (Louisiana and Georgia) using a different set of 30 images from these populations because these wasps tend to have darker reds than those in more northerly populations. Finally, we classified the pixels of all images to the nearest of these three colors in the palettes using the ‘imposeColors’ function in *recolorize*.

To quantify variation among individuals, we converted the images back to rasters consisting of a stack of three binary rasters corresponding to pixel assignments for each of the three colors. Because we were interested in pattern variation, we treated the slightly different black and red colors of the northern and southern wasps as equivalent. We then analyzed variation using *patternize* and computed a principal components analysis of these rasters which yielded 269 components corresponding to the 269 images in the data set. We reduced this dataset to 24 statistically significant components (Fig. S7, Table S2), which were determined using permutation parallel analysis in the *jackstraw* package^102^. We then computed pairwise Euclidian distances between points in this multi-dimensional PCA space and quantified within-site face diversity as the mean pairwise distance between points collected from the same site.

### Cooperative nesting data

We obtained data on the number of foundresses per nest across the latitudinal range of this species using a combination of existing datasets compiled in WASPnest^31,43^ and our own observations of nesting behavior. For the WASPnest dataset, we restricted the dataset to observations where the number of foundresses was directly reported. We also excluded observations where the exact number of foundresses were unclear, for example if a paper simply stated that nests were “multi-foundress” without providing the number. We supplemented this dataset with our own observations of foundress associations across the range, including in some key populations at the southern end of the range. We observed nests early in the season before workers emerged. We also observed nests early in the morning or on cool and rainy days when all individuals associated with a nest tend to be on the nest. In total, this dataset consisted of 2,021 nest observations. We statistically analyzed the relationship between the number of foundresses per nest and latitude using a zero-truncated Poisson regression using the *VGAM* package^103^.

### Genomic analyses

To confirm that northern and southern *P. fuscatus* were the same species, we collected and sequenced the genomes of unrelated female *P. fuscatus* from five populations: New York (*n* = 30), Massachusetts (*n* = 10), North Carolina (*n* = 8), Georgia (*n* = 15), and Louisiana (*n* = 25). As an outgroup, we included three individuals each from three closely related species (*P. carolina*, *P. dorsalis*, and *P. metricus*) with sympatric ranges. Sample information is provided in Table S4.

Paired-end 150-bp Nextera libraries were sequenced on the Illumina HiSeq 2000. All samples were aligned to the *P. fuscatus* reference genome^52^ using the Burrow-Wheeler Aligner (v.0.7.13)^104^. Variants were identified using GATK (v3.8)^105^ and hard filtered to remove low confidence variants, following the methods described in^45^.

To examine the relationship between samples, we constructed a phylogenetic tree with SNPhylo (v20160204)^106^, a program designed to rapidly build phylogenetic trees from large SNP datasets. To reduce the size of the dataset, variants were first filtered with VCFtools^107^ to retain only a single, informative, high-quality, biallelic SNP every 1,000 bp using the options: --max-alleles 2 --mac 0.1 --max-missing-count 10 --min-meanDP 3 --max-meanDP 1200 --minQ 20 --thin 1000. SNPhylo was run with 500 rounds of bootstrapping. We further explored relatedness between samples by conducting a PCA of genetic variants using Tassel5^108^. Lastly, we calculated genetic differentiation between the most distant populations, New York and Louisiana, using Weir-Cockerham FST, implemented in VCFtools.

### Recent selection in northern versus southern wasps

Using the 40 re-sequenced *P. fuscatus* genomes from Georgia and Louisiana, we looked for evidence of selective sweeps in southern wasps with SweepFinder2^53^. SweepFinder2 uses deviations in the local site frequency spectrum to infer selective sweeps, generating a composite likelihood ratio (CLR) value for each window. CLR values are larger when selection is stronger, more recent, and/or acting on new mutations rather than standing genetic variation ^52^. We compared CLR values for the southern population to CLR values that were generated for a prior study of northern populations^52^. Northern CLR values were calculated from the same 40 wasps from New York and Massachusetts described above. We included two sampling sites in each analysis to avoid detecting selective sweeps caused by local adaptation. CLR values between northern and southern wasps were scaled by the maximum CLR value in each dataset, generating scaled CLR. Values were compared in 1000 bp windows across the genome and plots were constructed with BioCircos^109^. For each gene in the genome, as well as the region +/- 5000 bp upstream/downstream of that gene, we calculated a maximum scaled CLR value.

Genes in the *P. fuscatus* genome had been previously classified as potential targets of selection for cognitive evolution if annotated with one of the following Gene Ontology (GO) terms: cognition (GO:0050890), mushroom body development (GO:0016319), visual behavior (GO:0007632), learning or memory (GO:0007611), and eye development (GO:0001654). Out of 11,935 genes, 1,088 genes were considered potentially related to the perceptual and cognitive mechanisms of individual recognition (hereafter: ‘visual cognition genes’). We also categorized genes based on whether or not they showed evidence of differential expression in response to social experience based on data published in^54^. For both data sets, to statistically compare scaled CLR values between populations and gene categories, we log transformed scaled CLR values to improve linearity and fit linear mixed effects models using the *lme4* package, with population (northern or southern), gene type (GO term dataset: visual cognition gene or other; differential expression dataset: yes or no), and their interaction as fixed effects, and gene identity as a random effect. We evaluated the significance of fixed effects and their interaction using type III ANOVAs using the *car* package, and we report Wald chi-square test statistics. We visualized population-specific elevation of CLR values for candidate social cognition loci by computing the residual CLR value per locus. To do this, we generated expected CLR values by randomly selecting 100 sets of *n* non-candidate loci, where *n* is the number of candidate loci for a dataset, i.e., *n* = 1,088 genes based on visual cognition GO terms, *n* = 733 genes for socially regulated genes. We then ranked each set by decreasing CLR value and took the mean CLR value at each rank across the 100 sets to estimate expected CLR values for *n* random loci^110^. We also ranked the observed CLR values for candidate loci and took the difference between the observed CLR value and expected CLR value for each rank as the residual CLR. These residuals thus control for potential population differences in CLR values across the genome and allow visualization of potential differences in the elevation of CLR values for candidate loci.

## Data accessibility

New sequence data for samples from Louisiana and Georgia have been deposited to the NCBI sequence read archive in project PRJNA761367. Samples from other populations are available in project PRJNA482994. All other data will be made publicly available on Dryad upon publication.

## Acknowledgements

We thank the state parks departments of New York, Pennsylvania, Virginia, North Carolina, South Carolina, Georgia, Alabama, and Louisiana for permission to collect paper wasps. We thank Benjamin Brack for assistance in setting up behavioral experiments and Lilly Woodward for help processing photographs. This research was funded by a National Science Foundation CAREER grant to MJS (DEB-1750394) and a grant from the National Institutes of Health to MJS (DP2-GM128202). AW was supported by an NSF EDGE grant to A. Toth and MJS (1827567).

## Author contributions

JPT and MJS conceived of and designed the project; JPT and CMJ designed and performed the individual recognition experiment; JPT performed other behavioral experiments; JPT, CMJ, SV, RS, AWL, TJP, FMKU, and AW collected data; SMVB and HIW developed code for color pattern analysis; JPT analyzed non-genomic data; SEM and MJS analyzed genomic data; MJS secured funding; JPT, SEM, and MJS wrote the first draft of the paper; and all authors reviewed and edited the paper.

