## Supplementary Material for "Evidence for a selective link between cooperation and individual recognition"

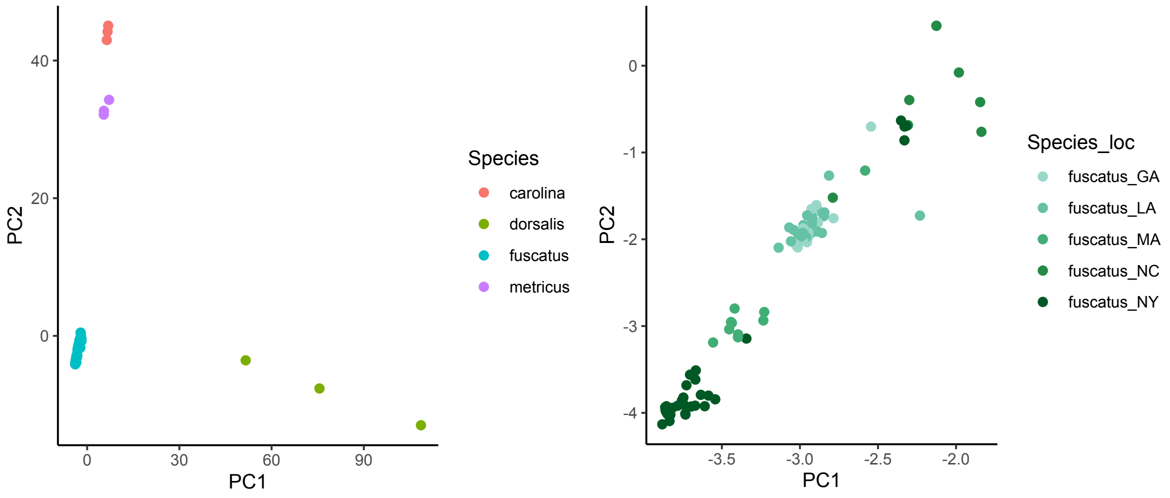

**Figure S1.** The first two principal components from a PCA of SNP data from aligned genomic data. On the left, samples from *Polistes fuscatus* are shown along with three other closely related species (*P. carolina*, *P. dorsalis*, and *P. metricus*). On the right, we visualize a PCA of SNP data for *P. fuscatus* samples alone, with different colors designating different US states where the wasps were collected.

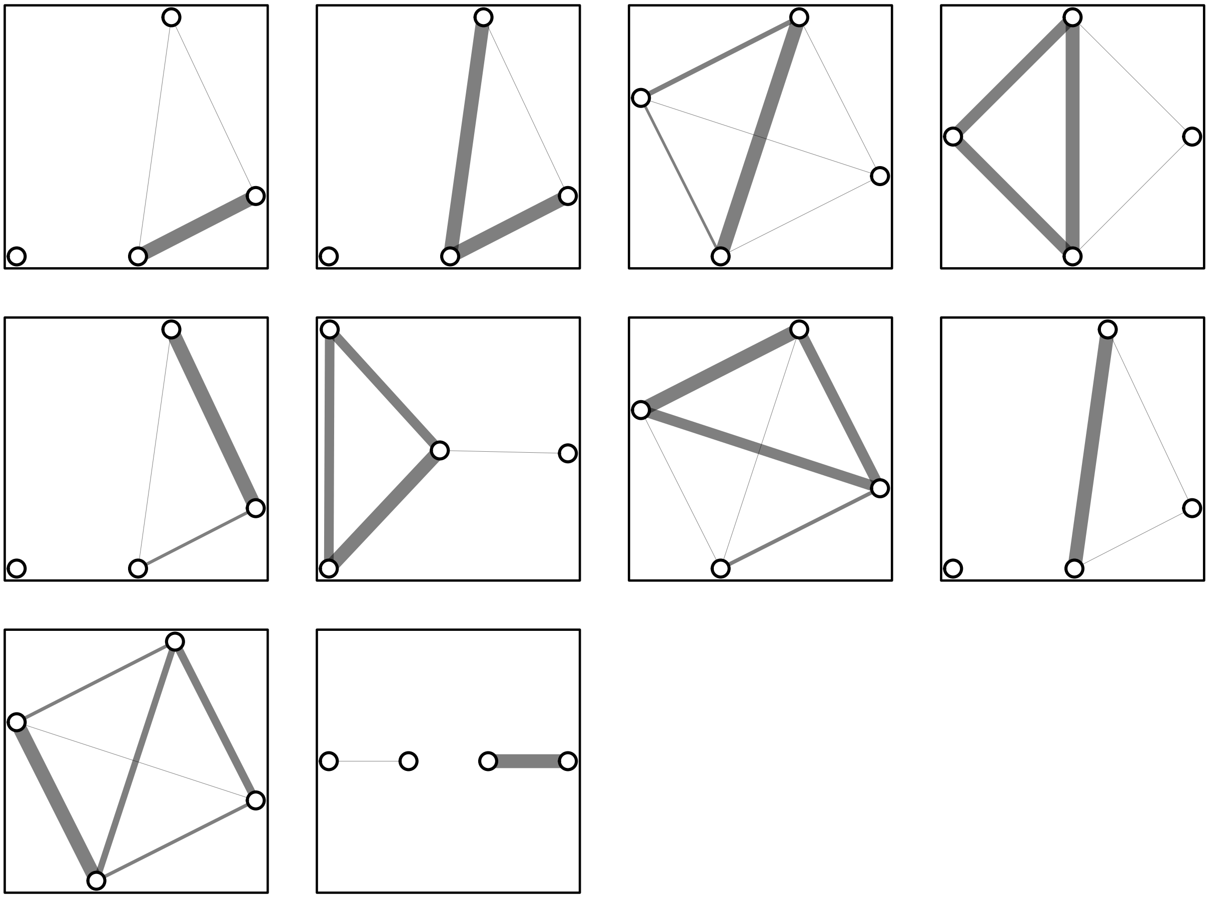

**Figure S2.** Social networks of all 10 northern groups of wasps, derived from nocturnal “huddle” associations of wasps.

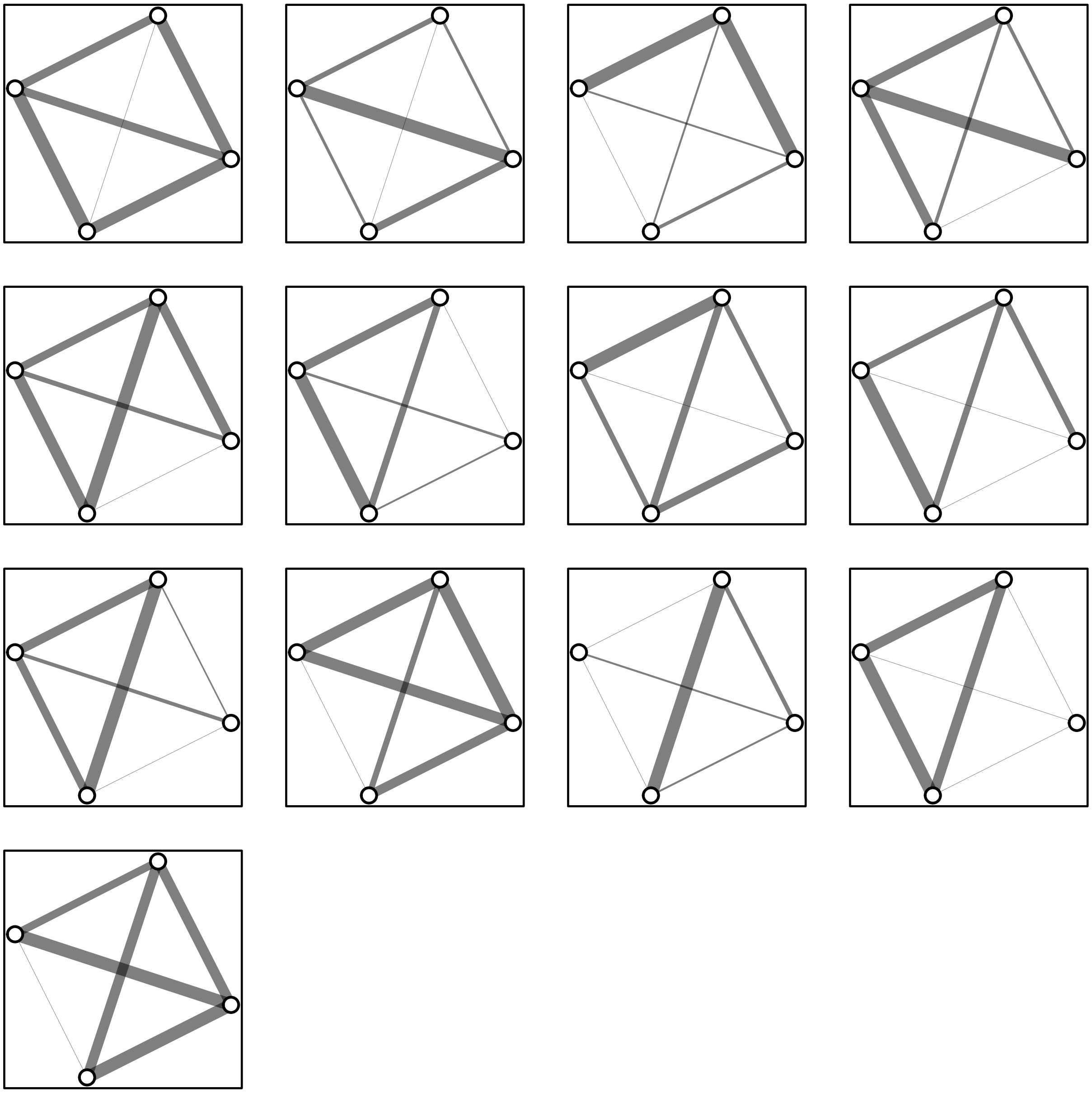

**Figure S3.** Social networks of all 13 groups of southern wasps, derived from nocturnal “huddle” associations of wasps.

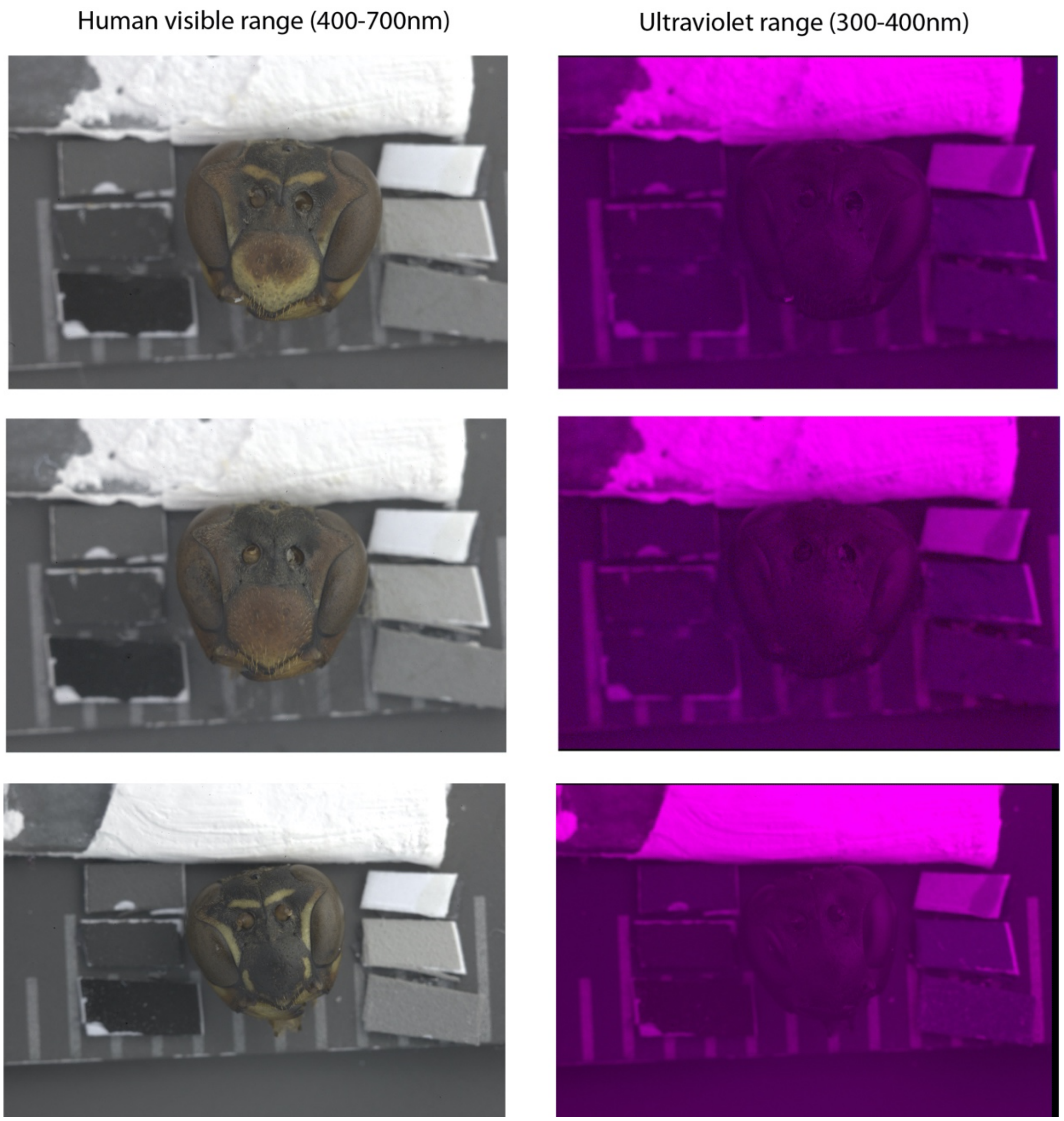

**Figure S4.** Comparison of three *Polistes fuscatus* faces in the human visual range (400-700nm) and the ultraviolet range (300-400nm). Photographs were taken using a converted Canon 6D camera with the camera UV filter removed and a Nikon 80mm f.5.6 EL-Nikkor lens. The visual range photographs were taken using a Hoya UV and IR cut filter to block light from outside of the visible spectrum, while the UV photographs were taken using a Baader Planetarium U-filter 2, which allows imaging from 300-400nm in the UV spectrum. Visible range photographs were illuminated with compact fluorescent lights (see illumination spectrum in Fig. S5) and UV photographs were illuminated with Exo Terra SunRay lamp equipped with a 35W Sunray bulb. In each photograph are a series of 6 spectrally flat gray standards (DGK Color Tools) that reflect human visual light, as well as a UV white standard made from white reflective paint with reflectance values of 95 to 98% from 300 to 1200 nm (Labsphere 6080 white reflectance coating). Photographs were taken in RAW format, and then normalized with respect to lighting and combined to a multispectral image using the mica Toolbox (Troscianko & Stevens 2015). We then visualized the visual range of these images (left) by creating a presentation image of the red, green, and blue color channels. We visualized the UV range of the images by creating a presentation image and coloring the UV-R channel with red, and the UV-B channel with blue. The reflectance on the UV-sensitive white standard on top confirms that these images are reflecting UV light, and the absence of UV reflectance seen on the wasp faces confirms that the colors on their faces do not reflect light in the UV range.

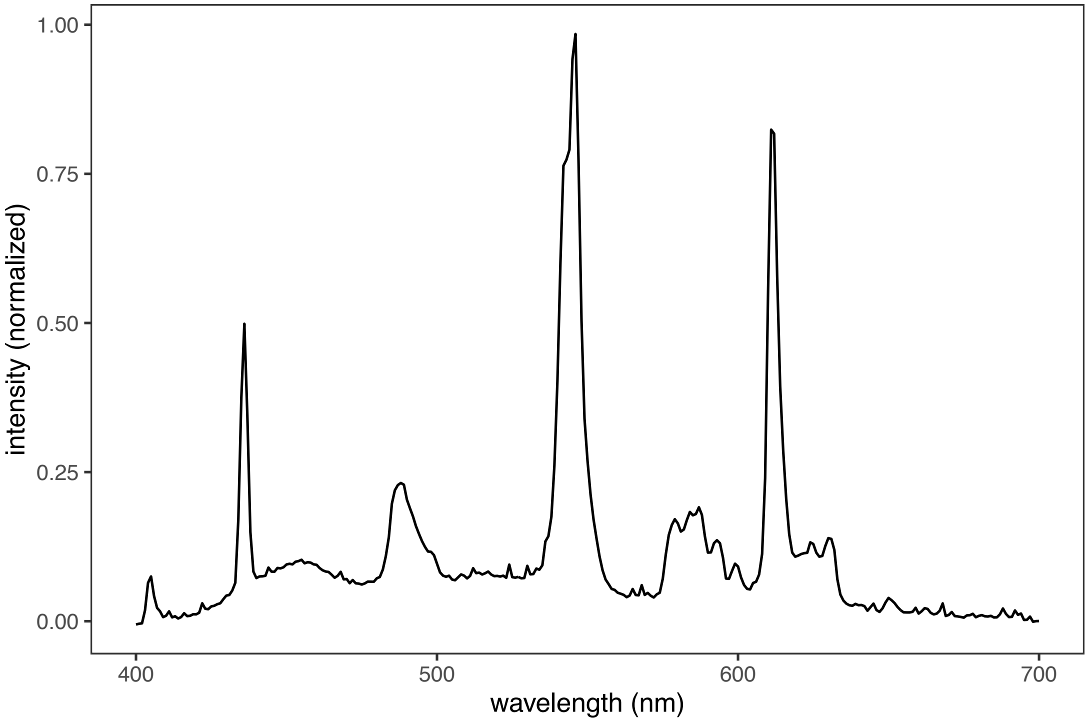

**Figure S5**. Illumination spectrum for human-visible range (400-700nm) wasp photography. The spectrum was measured with an OceanOptics SD 2000 spectrometer (OceanOptic, Dunedin, FL, U.S.A), with the spectrometer positioned facing upwards at the exact location of where the wasp specimens are placed for photography. Measurement was taken using 2,000ms integration time, 1nm boxcar width and averaged across 3 scans.

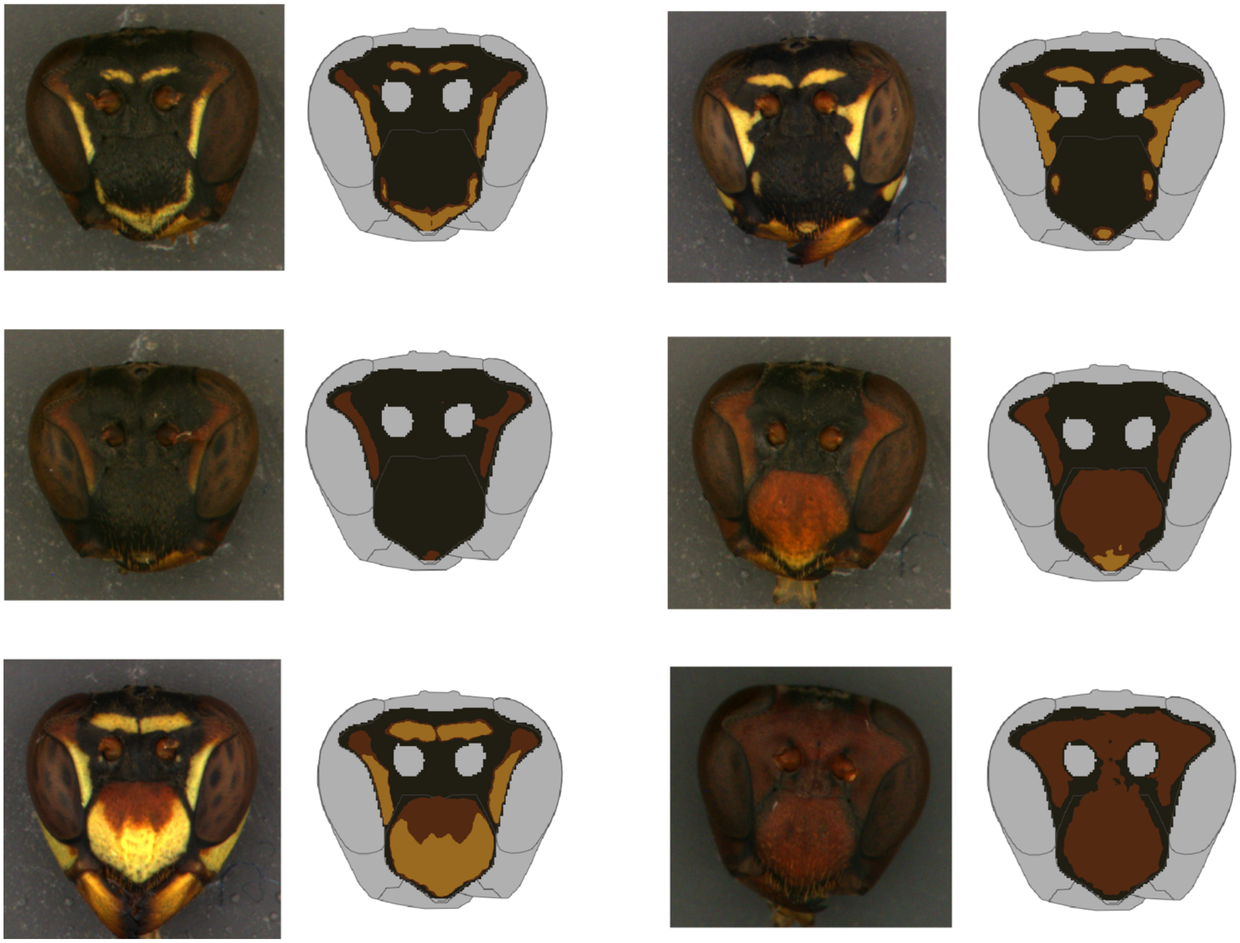

**Fig. S6.** Example face photographs (left columns) and corresponding color segmented face images (left columns). For color segmentation, areas outside of regions of interest (e.g., eyes, mandibles, antennae), were masked, and then colors were classified to the nearest of three main colors: yellow, red/brown, black (see Methods for details).

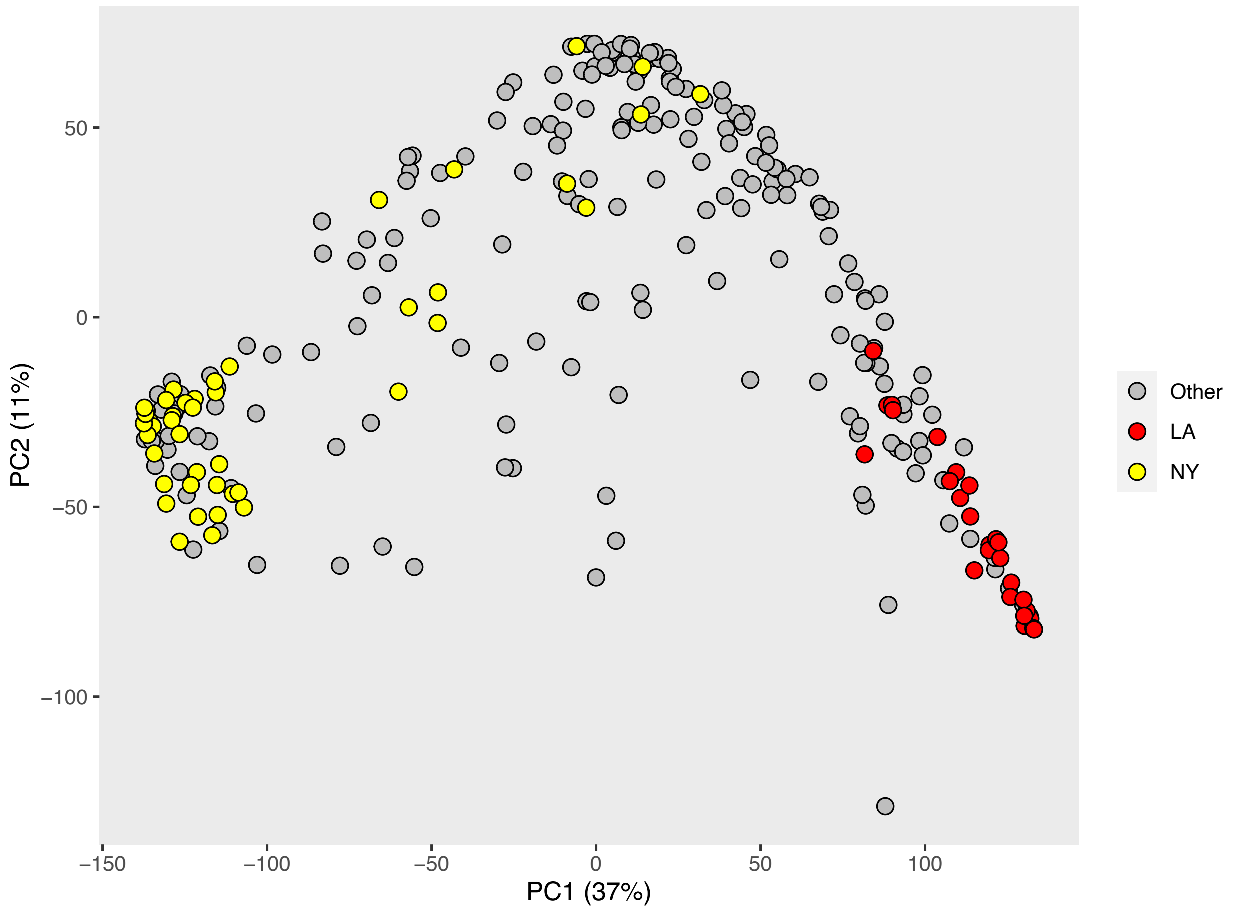

**Figure S7.** The first two principal components of the color pattern PCA, with wasps from New York (yellow) and Louisiana (red) highlighted.

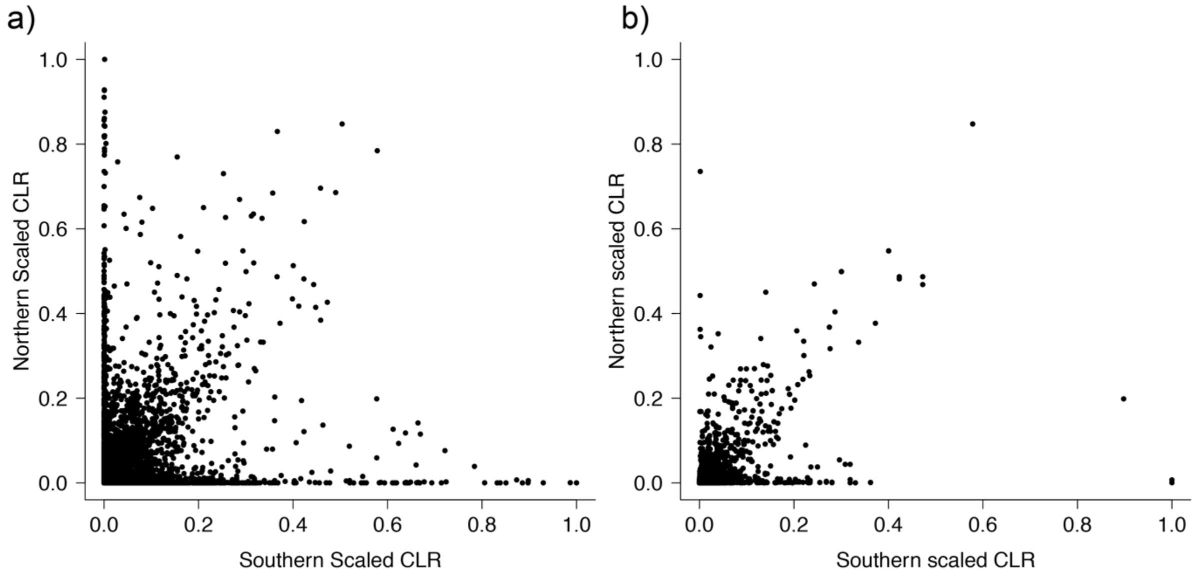

**Figure S8. Shared and independent regions under selection between northern and southern populations.** a) Genome-wide comparison of scaled composite likelihood ratio (CLR) values between northern and southern populations. Each point shows mean CLR for a 1,000 bp window. b) Comparison of CLR values between populations at each gene in the *P. fuscatus* genome. Each point is the maximum CLR value of a gene including the region +/- 5000 bp upstream/downstream of the coding sequence.

**Table S1.** Factor loadings of each aggressive behavior on first principal component of a PCA, which was used as an aggression index in the individual recognition experiment.

| Behavior | Aggression Index (PC1) |
|---|---|
| Approach | 0.205 |
| Bite | 0.463 |
| Dart | 0.207 |
| Dodge | 0.350 |
| Kick | 0.331 |
| Snap | 0.500 |
| Antennation | 0.444 |
| Chase | 0.136 |
| Grapple | 0.052 |
| Standard deviation | 1.721 |
| Proportion of Variance | 0.329 |

**Table S2.** Summary of importance of each principal component from the PCA analysis of color segmented face images. The first 24 PCs were statistically significant.

| | PC1 | PC2 | PC3 | PC4 | PC5 | PC6 | PC7 | PC8 | PC9 | PC10 | PC11 | PC12 |
|---|---|---|---|---|---|---|---|---|---|---|---|---|
| Standard deviation | 83.5533769 | 45.9083657 | 39.7420807 | 23.7186718 | 21.7601505 | 20.8136458 | 18.018592 | 16.6042699 | 14.9028918 | 14.0659496 | 13.523655 | 12.3650433 |
| Proportion of Variance | 0.36604 | 0.11051 | 0.08281 | 0.0295 | 0.02483 | 0.02271 | 0.01702 | 0.01446 | 0.01165 | 0.01037 | 0.00959 | 0.00802 |
| Cumulative Proportion | 0.36604 | 0.47655 | 0.55936 | 0.58886 | 0.61369 | 0.6364 | 0.65343 | 0.66788 | 0.67953 | 0.6899 | 0.69949 | 0.70751 |
| | PC13 | PC14 | PC15 | PC16 | PC17 | PC18 | PC19 | PC20 | PC21 | PC22 | PC23 | PC24 |
| Standard deviation | 11.743583 | 11.2304747 | 10.5713084 | 10.3658838 | 10.2729844 | 9.94302721 | 9.82686772 | 9.58659645 | 9.05565315 | 8.88992105 | 8.77988777 | 8.60993782 |
| Proportion of Variance | 0.00723 | 0.00661 | 0.00586 | 0.00563 | 0.00553 | 0.00518 | 0.00506 | 0.00482 | 0.0043 | 0.00414 | 0.00404 | 0.00389 |
| Cumulative Proportion | 0.71474 | 0.72135 | 0.72721 | 0.73285 | 0.73838 | 0.74356 | 0.74863 | 0.75345 | 0.75775 | 0.76189 | 0.76593 | 0.76982 |

**Table S3.** The numbers of visual cognition genes and genes differentially expressed genes in response to social interactions, as a percentage of the total number of genes, in the whole genome and among the top 5% of CLR scores. In northern populations, visual cognition genes and social differentially expressed genes make up a disproportionately high percentage of genes with the highest CLR values, and these percentages are higher than in southern populations.

| | Whole genome | Top 5% CLR scores |
|---|---|---|
| *Visual cognition genes* | | |
| North | 9% (1088/11935) | 14% (83/598) |
| South | 9% (1088/11935) | 10% (57/597) |
| *Social differentially expressed genes* | | |
| North | 6% (733/11652) | 14% (80/585) |
| South | 6% (733/11652) | 9% (55/582) |

**Table S4.** Sample information for the 97 re-sequenced genomes used in this study. Raw data has been archived at the NCBI Sequence Read Archive under Bioproject accession numbers PRJNA482994 and PRJNA761367.

| Species | Sample ID | Region | Location | Latitude | Longitude | SRA ID |
|---|---|---|---|---|---|---|
| carolina | Carolina_AR_H197 | AR | Arkadelphia | 34.12522 | -93.05990 | SAMN09728973 |
| carolina | Carolina_AR_H201 | AR | Arkadelphia | 34.12522 | -93.05990 | SAMN09728974 |
| carolina | Carolina_TN_H223 | TN | Lebanon | 36.09172 | -86.33080 | SAMN09728976 |
| dorsalis | Dorsalis_NY_474 | NY | Newfield | 42.53319 | -76.31308 | SAMN09728924 |
| dorsalis | Dorsalis_NY_495 | NY | Elmira | 42.09551 | -76.81706 | SAMN09728925 |
| dorsalis | Dorsalis_NY_EHCC7 | NY | Ithaca | 42.43050 | -76.41417 | SAMN09728929 |
| fuscatus | JPT_2019_253 | GA | Jekyll Island | 31.30444 | -81.59889 | |
| fuscatus | JPT_2019_254 | GA | Jekyll Island | 31.30444 | -81.59889 | |
| fuscatus | JPT_2019_272 | GA | Brunswick | 31.15444 | -81.55472 | |
| fuscatus | JPT_2019_287 | GA | Brunswick | 31.28444 | -81.58806 | |
| fuscatus | JPT_2019_288 | GA | Brunswick | 31.28444 | -81.58806 | |
| fuscatus | JPT_2019_289 | GA | Brunswick | 31.28444 | -81.58806 | |
| fuscatus | JPT_2019_290 | GA | Brunswick | 31.28444 | -81.58806 | |
| fuscatus | JPT_2019_291 | GA | Brunswick | 31.28444 | -81.58806 | |
| fuscatus | JPT_2019_301 | GA | Skidaway Institute of Oceanography | 32.08472 | -81.08722 | |
| fuscatus | JPT_2019_302 | GA | Skidaway Institute of Oceanography | 32.08472 | -81.08722 | |
| fuscatus | JPT_2019_338 | GA | Skidaway Institute of Oceanography | 32.07778 | -81.08250 | |
| fuscatus | JPT_2019_381 | GA | Skidaway Institute of Oceanography | 32.05056 | -81.11667 | |
| fuscatus | JPT_2019_383 | GA | Skidaway Institute of Oceanography | 32.05056 | -81.11667 | |
| fuscatus | JPT_2019_385 | GA | Skidaway Institute of Oceanography | 32.05056 | -81.11667 | |
| fuscatus | JPT_2019_388 | GA | Skidaway Institute of Oceanography | 32.05056 | -81.11667 | |
| fuscatus | JPT_2019_11 | LA | Fontainebleau State Park | 30.34361 | -90.04361 | |
| fuscatus | JPT_2019_112 | LA | Bogue Chitto State Park | 30.84139 | -90.36528 | |
| fuscatus | JPT_2019_114 | LA | Bogue Chitto State Park | 30.87556 | -90.37833 | |
| fuscatus | JPT_2019_116 | LA | Bogue Chitto State Park | 30.87556 | -90.37833 | |
| fuscatus | JPT_2019_119 | LA | Bogue Chitto State Park | 30.87556 | -90.37833 | |
| fuscatus | JPT_2019_12 | LA | Fontainebleau State Park | 30.34361 | -90.04361 | |
| fuscatus | JPT_2019_126 | LA | Bogue Chitto State Park | 30.97778 | -90.29250 | |
| fuscatus | JPT_2019_127 | LA | Bogue Chitto State Park | 30.97778 | -90.29250 | |
| fuscatus | JPT_2019_13 | LA | Fontainebleau State Park | 30.34361 | -90.04361 | |
| fuscatus | JPT_2019_137 | LA | Bogue Chitto State Park | 30.99194 | -90.37333 | |
| fuscatus | JPT_2019_140 | LA | Bogue Chitto State Park | 30.99194 | -90.37333 | |
| fuscatus | JPT_2019_18 | LA | Fontainebleau State Park | 30.34556 | -90.04083 | |
| fuscatus | JPT_2019_19 | LA | Fontainebleau State Park | 30.34556 | -90.04083 | |
| fuscatus | JPT_2019_20 | LA | Fontainebleau State Park | 30.34556 | -90.04083 | |
| fuscatus | JPT_2019_22 | LA | Fontainebleau State Park | 30.34556 | -90.04083 | |
| fuscatus | JPT_2019_3 | LA | Fontainebleau State Park | 30.34361 | -90.04361 | |
| fuscatus | JPT_2019_4 | LA | Fontainebleau State Park | 30.34361 | -90.04361 | |
| fuscatus | JPT_2019_6 | LA | Fontainebleau State Park | 30.34361 | -90.04361 | |
| fuscatus | JPT_2019_66 | LA | Palmetto Island State Park | 30.09056 | -92.40194 | |
| fuscatus | JPT_2019_7 | LA | Fontainebleau State Park | 30.34361 | -90.04361 | |
| fuscatus | JPT_2019_74 | LA | Palmetto Island State Park | 30.00333 | -92.29056 | |
| fuscatus | JPT_2019_75 | LA | Palmetto Island State Park | 30.00333 | -92.29056 | |
| fuscatus | JPT_2019_76 | LA | Palmetto Island State Park | 30.00333 | -92.29056 | |
| fuscatus | JPT_2019_8 | LA | Fontainebleau State Park | 30.34361 | -90.04361 | |
| fuscatus | JPT_2019_9 | LA | Fontainebleau State Park | 30.34361 | -90.04361 | |
| fuscatus | Fuscatus_MA_362 | MA | Carver | 41.87367 | -70.72940 | SAMN09728903 |
| fuscatus | Fuscatus_MA_365 | MA | East Freetown | 41.77944 | -70.92139 | SAMN09728904 |
| fuscatus | Fuscatus_MA_368 | MA | Freetown | 41.76715 | -70.93685 | SAMN09728905 |
| fuscatus | Fuscatus_MA_373 | MA | Freetown | 41.76715 | -70.93685 | SAMN09728906 |
| fuscatus | Fuscatus_MA_375 | MA | East Freetown | 41.77944 | -70.92139 | SAMN09728907 |
| fuscatus | Fuscatus_MA_377 | MA | Middleborough | 41.82583 | -70.83528 | SAMN09728908 |
| fuscatus | Fuscatus_MA_383 | MA | Carver | 41.87361 | -70.72694 | SAMN09728909 |
| fuscatus | Fuscatus_MA_386 | MA | Rochester | 41.75189 | -70.88180 | SAMN09728910 |

| | | | | | | |
|---|---|---|---|---|---|---|
| fuscatus | Fuscatus_MA_391 | MA | Middleborough | 41.84417 | -70.84833 | SAMN09728911 |
| fuscatus | Fuscatus_MA_393 | MA | Middleborough | 41.80750 | -70.88028 | SAMN09728912 |
| fuscatus | Fuscatus_NC_2016_07 | NC | Raleigh | 36.00393 | -78.92310 | |
| fuscatus | Fuscatus_NC_2016_08 | NC | Raleigh | 36.00393 | -78.92310 | SAMN10485636 |
| fuscatus | Fuscatus_NC_2016_13 | NC | Raleigh | 36.01250 | -78.90160 | SAMN10485637 |
| fuscatus | Fuscatus_NC_2016_17 | NC | Raleigh | 36.00105 | -78.96170 | SAMN10485638 |
| fuscatus | Fuscatus_NC_2017_09 | NC | Raleigh | 36.01836 | -78.89720 | |
| fuscatus | Fuscatus_NC_2017_10 | NC | Raleigh | 36.01836 | -78.89720 | SAMN10485639 |
| fuscatus | Fuscatus_NC_2017_11 | NC | Raleigh | 36.01836 | -78.89720 | SAMN10485640 |
| fuscatus | Fuscatus_NC_2017_12 | NC | Raleigh | 36.01836 | -78.89720 | SAMN10485641 |
| fuscatus | Fuscatus_NY_10 | NY | Freeville | 42.53319 | -76.31308 | SAMN09728875 |
| fuscatus | Fuscatus_NY_14 | NY | Arnot Forest | 42.26503 | -76.62827 | SAMN09728876 |
| fuscatus | Fuscatus_NY_20 | NY | Spencer | 42.34694 | -76.48723 | SAMN09728877 |
| fuscatus | Fuscatus_NY_22 | NY | Arnot Forest | 42.26514 | -76.62831 | SAMN09728878 |
| fuscatus | Fuscatus_NY_401 | NY | Slaterville Springs | 42.39435 | -76.34641 | SAMN09728879 |
| fuscatus | Fuscatus_NY_402 | NY | Slaterville Springs | 42.39435 | -76.34641 | SAMN09728880 |
| fuscatus | Fuscatus_NY_431 | NY | Ithaca | 42.45172 | -76.47994 | SAMN09728881 |
| fuscatus | Fuscatus_NY_525 | NY | Arnot Forest | 42.26503 | -76.62827 | SAMN09728882 |
| fuscatus | Fuscatus_NY_526 | NY | Arnot Forest | 42.26503 | -76.62827 | SAMN09728883 |
| fuscatus | Fuscatus_NY_528 | NY | Arnot Forest | 42.26503 | -76.62827 | SAMN09728884 |
| fuscatus | Fuscatus_NY_533 | NY | Arnot Forest | 42.26503 | -76.62827 | SAMN09728885 |
| fuscatus | Fuscatus_NY_540 | NY | Arnot Forest | 42.26503 | -76.62827 | SAMN09728886 |
| fuscatus | Fuscatus_NY_545 | NY | Arnot Forest | 42.26503 | -76.62827 | SAMN09728887 |
| fuscatus | Fuscatus_NY_550 | NY | Arnot Forest | 42.26503 | -76.62827 | SAMN09728888 |
| fuscatus | Fuscatus_NY_552 | NY | Arnot Forest | 42.26503 | -76.62827 | SAMN09728889 |
| fuscatus | Fuscatus_NY_560 | NY | Arnot Forest | 42.26503 | -76.62827 | SAMN09728890 |
| fuscatus | Fuscatus_NY_568 | NY | Arnot Forest | 42.26503 | -76.62827 | SAMN09728891 |
| fuscatus | Fuscatus_NY_575 | NY | Arnot Forest | 42.26503 | -76.62827 | SAMN09728892 |
| fuscatus | Fuscatus_NY_576 | NY | Arnot Forest | 42.26503 | -76.62827 | SAMN09728893 |
| fuscatus | Fuscatus_NY_582 | NY | Arnot Forest | 42.26503 | -76.62827 | SAMN09728894 |
| fuscatus | Fuscatus_NY_585 | NY | Arnot Forest | 42.26503 | -76.62827 | SAMN09728895 |
| fuscatus | Fuscatus_NY_590 | NY | Arnot Forest | 42.26503 | -76.62827 | SAMN09728896 |
| fuscatus | Fuscatus_NY_620 | NY | Spencer | 42.20780 | -76.48517 | SAMN09728897 |
| fuscatus | Fuscatus_NY_628 | NY | Erin | 42.18059 | -76.68637 | SAMN09728898 |
| fuscatus | Fuscatus_NY_638 | NY | Freeville | 42.53319 | -76.31308 | SAMN09728899 |
| fuscatus | Fuscatus_NY_642 | NY | Freeville | 42.53319 | -76.31308 | SAMN09728900 |
| fuscatus | Fuscatus_NY_647 | NY | Freeville | 42.53319 | -76.31308 | SAMN09728901 |
| fuscatus | Fuscatus_NY_658 | NY | Freeville | 42.53319 | -76.31308 | SAMN09728902 |
| fuscatus | Fuscatus_NY_7 | NY | Arnot Forest | 42.26503 | -76.62827 | SAMN09728873 |
| fuscatus | Fuscatus_NY_9 | NY | Freeville | 42.53319 | -76.31308 | SAMN09728874 |
| metricus | Metricus_MD_17_6 | MD | Marriottsville | 39.35347 | -76.88737 | SAMN09728934 |
| metricus | Metricus_NC_31 | NC | Raleigh | 36.01250 | -78.90163 | SAMN09728935 |
| metricus | Metricus_NC_32 | NC | Raleigh | 36.01250 | -78.90163 | SAMN09728936 |
